# CASP16 protein monomer structure prediction assessment

**DOI:** 10.1101/2025.05.29.656942

**Authors:** Rongqing Yuan, Jing Zhang, Andriy Kryshtafovych, R. Dustin Schaeffer, Jian Zhou, Qian Cong, Nick V. Grishin

**Affiliations:** Eugene McDermott Center for Human Growth and Development, University of Texas Southwestern Medical Center, Dallas, TX, USA; Department of Biophysics, University of Texas Southwestern Medical Center, Dallas, TX, USA; Lyda Hill Department of Bioinformatics, University of Texas Southwestern Medical Center, Dallas, TX, USA; Genome Center, University of California, Davis, California, USA; Harold C. Simmons Comprehensive Cancer Center, University of Texas Southwestern Medical Center, Dallas, TX, USA; Department of Biochemistry, University of Texas Southwestern Medical Center, Dallas, TX, USA

**Keywords:** CASP16, protein structure prediction, monomer, AlphaFold2, AlphaFold3, quality estimation, stoichiometry, structure sampling

## Abstract

The assessment of monomer targets in the Critical Assessment of Structure Prediction Round 16 (CASP16) underscores that the problem of single-domain protein fold prediction is nearly solved—no target folds were incorrectly predicted across all Evaluation Units. However, challenges remain in accurately modeling truncated sequences, irregular secondary structures, and interaction-induced conformational changes. The release of AlphaFold3 (AF3) during CASP16, and its effective integration by many groups, demonstrated its superiority over AlphaFold2 (AF2), particularly in confidence estimation and model selection. Additional improvements in multiple sequence alignments (MSAs) and fragment-based prediction, i.e., selecting the optimal fragment of the full sequence for modeling, also contributed to enhanced prediction accuracy. The top three groups—all from the Yang lab—consistently outperformed others across CASP16 monomer targets, reflecting their robust modeling pipelines and successful adoption of AF3. CASP16 also introduced three new challenges: Phase 0, in which stoichiometry was withheld; Phase 2, which supplied ∼8,000 MassiveFold models per target to test model selection strategies; and Model 6, which limited predictors to using MSAs provided by the organizers. While we evaluated group performance in these additional challenges, the insights gained were limited due to low participation and caveats in the design of experiments. We suggest improvements for the organization of these challenges and encourage broader engagement from the prediction community. The progress in monomer modeling from CASP15 to CASP16 was subtle, but more groups in CASP16 were able to outperform ColabFold, reflecting the community’s improved ability in optimizing AF2 and the growing adoption of AF3. We anticipate that the recent release of the AF3 source code will stimulate future progress through user-driven optimization and innovations in model architecture. Finally, model ranking remains a persistent weakness across most groups, highlighting a critical area for future development.

## Introduction

Starting from 1994, the Critical Assessment of Structure Prediction (CASP) experiments^1^, conducted biennially, have played a pivotal role in establishing a community-wide standard for protein structure prediction. CASP evaluates the performance of emerging computational methods, identifies the promising trends, and pushes the community’s progress forward. The design of CASP experiments is as follows. Before the protein structures—determined through X-ray crystallography, cryo-electron microscopy, or nuclear magnetic resonance (NMR)—are publicly released and deposited into the Protein Data Bank (PDB)^2,3^, their information is submitted to the CASP organizers. The organizers release suitable sequences for prediction. Participating groups submit predicted protein 3D structures based on the sequences. Subsequently, independent assessors are invited to evaluate the performance blindly using standardized metrics and statistical models, ranking groups based on their accuracy. More importantly, the assessors will also highlight significant methodological advances and pinpoint areas in need of further development^4^.

The continuous progress in this field, in particular, the introduction of coevolution-based contact prediction and deep learning methods, has culminated in the breakthrough in protein structure prediction by AlphaFold2 (AF2) during CASP14^5–9^, which has stimulated numerous scientific discoveries and biomedical applications^10–22^. AF2 has become an essential component for subsequent method development among CASP predictors, and winning strategies in CASP15 all built on top of AF2^23–25^. At the beginning of the current CASP (CASP16), DeepMind released the AlphaFold3 (AF3) web server^26^, and multiple CASP16 predictors thus incorporated AF3 in their structure prediction pipeline. Assessing whether AF3 represents another significant improvement in the field is therefore a critical assessment task of CASP16.

This assessment focuses on monomer structure prediction, which has been at the core of the CASP experiment since its establishment^1^. However, since AF2 nearly solved the single-chain protein structure prediction problem in 2020, modeling monomer proteins has become relatively less challenging in recent years. Consequently, many monomer targets in this CASP are derived from homo-oligomeric protein complexes, multi-protein complexes, and protein-nucleotide complexes. Good performance in these monomer targets requires accurate modeling of the protein complexes, and thus, evaluation of monomer prediction accuracy also reflects CASP predictors’ ability in complex structure modeling.

To stimulate progress in the field, the following new challenges regarding protein and protein complex structure prediction were introduced in the current CASP:

**1. Phase 0.** In the traditional CASP routine (Phase 1 of CASP16), predictors are required to predict the 3D structure based on sequences and stoichiometry of a protein complex. CASP16 introduced Phase 0, where predictors were not provided with stoichiometry information. Predictors thus needed to solve the additional challenge of stoichiometry prediction, a factor significantly influencing the oligomeric structure prediction accuracy [cite: our oligomer evaluation paper], thereby affecting the monomer targets derived from protein complexes.
**2. Phase 2.** CASP16 also introduced Phase 2, in which predictors were provided with a large number (in most cases, 8,040 per target) of predicted structures by MassiveFold^27^. Massive sampling approaches using AlphaFold-like methods, exemplified by AFsample from the Wallner group, were a successful strategy in CASP15^28,29^. Thus, providing every predictor with large pools of predicted models is expected to stimulate progress in model quality estimation and model selection.
**3. Model 6.** Another winning strategy in CASP15 was improving upon the default multiple sequence alignments (MSAs) of AF2^30,23,31^. To examine the impact of deep learning network architecture independent of MSA quality, CASP16 introduced a special “model 6” aside from the traditional models 1–5. For model 6, predictors were asked to build structures using MSAs generated by ColabFold^32^. This additional challenge minimizes the influence of MSA quality and allows a focused evaluation of architectural innovations.

We herein report our assessment of the monomer prediction category in CASP16. In the following sections, we first analyze the strategies used by different groups and find that using AF3, enhancing MSAs, and refining the input query sequence (e.g., a fragment instead of the full sequence) for modeling were the winning strategies of CASP16. We then outline the features of the targets that influence the performance and highlight difficult targets. Finally, we provide the rankings of participants and our evaluations of the newly introduced CASP16 challenges. The results again demonstrate that protein monomer structure prediction is a largely solved problem^33^, and improvement in the monomer prediction accuracy in CASP16 compared to CASP15 is only subtle. We identify aspects that are still challenging to the field, such as predicting structures for truncated sequences, modeling irregular secondary structures, and selecting the best models. These areas represent directions for further advances by the community.

## Results and Discussions

### Overall view of targets and prediction strategies

For monomer categories, 125 human expert groups and 36 server groups submitted predictions for 16, 42, and 27 targets from Phase 0, 1, and 2, respectively. The Phase 0 monomer targets and models were all derived from Phase 0 oligomer targets and predictions submitted for these targets. These targets were defined from 11 monomeric proteins, 16 homo oligomeric protein complexes, 8 hetero oligomeric protein complexes, and 5 protein-nucleotide (hybrid) complexes. These proteins were first parsed into domains using DomainParser^34^, DDomain^35^, SWORD^36^, and Chainsaw^37^. Evaluation Units (EUs) were defined as single domains or joint domains if their relative orientations can be correctly predicted by most groups^38,39^. We defined 25, 74, and 46 EUs for Phase 0, 1, and 2 targets, respectively. We provided the definition of EUs for Phase 1 targets in the supplemental **Table. S1.**

An overview of the EUs, groups, and each group’s performance (measured by GDT_HA of the best model) is shown in **Fig. 1**. CASP encourages prediction of all targets to increase the statistical power of performance comparisons between groups, and the established evaluation routines penalize for missed targets. We evaluated the performance of all groups, but only those that submitted on more than a certain percentage of all EUs will be presented in the figures and tables, and the threshold will be denoted in the context. In **Fig. 1**, we only show groups that have predicted more than 25% (19) Phase 1 EUs.

**Figure 1.**
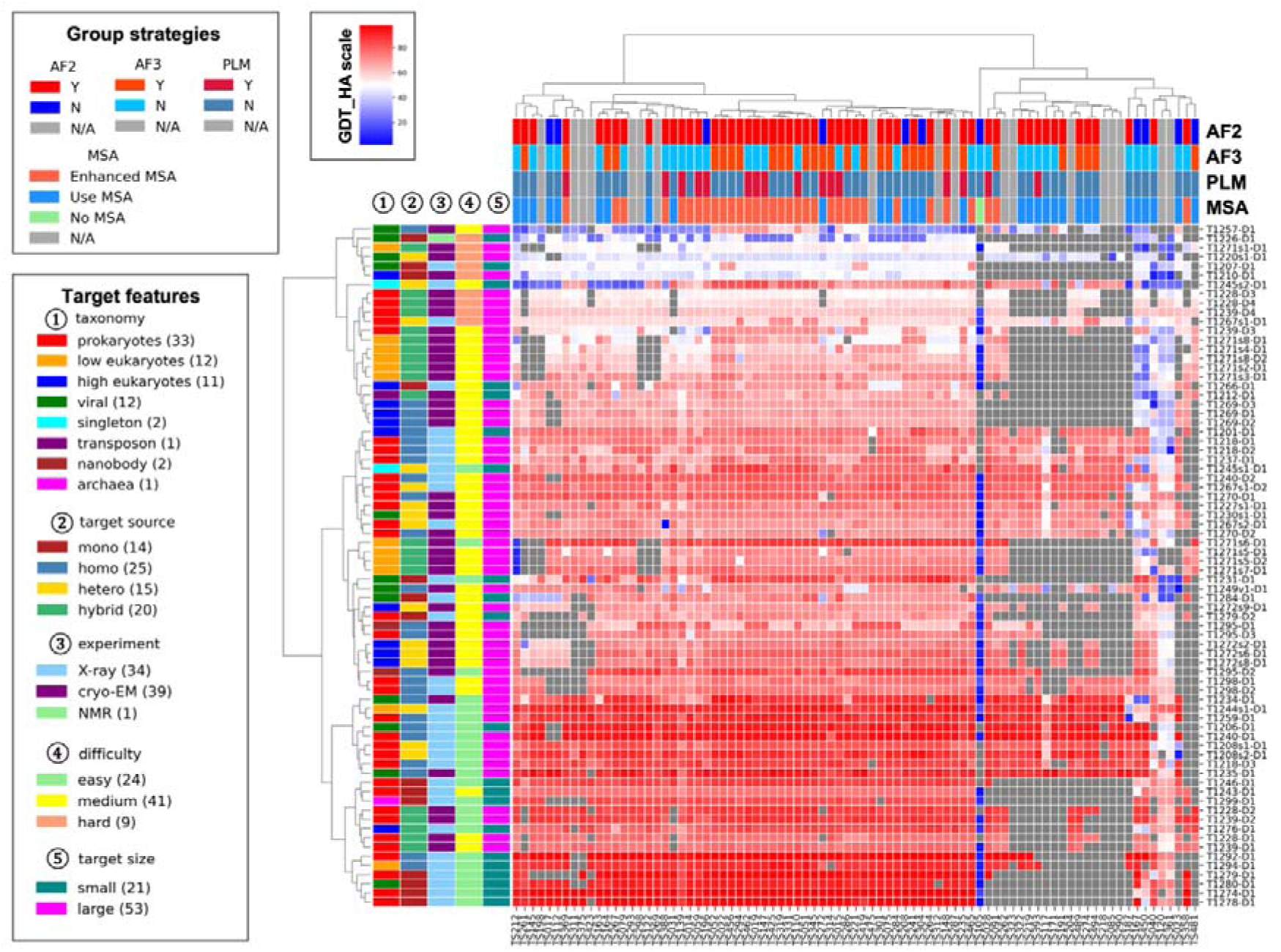
A heatmap of the targets and participating groups. The value in each box is the highest GDT_HA value of one group for one target (Grey indicates that such score is not available). The annotations on the x-axis and y-axis represent the different features of groups and targets, respectively. Only groups that submitted more than 25% of the models are included.

For most EUs, nearly all groups could generate high-quality models, and the performance differences are minimal. Some EUs turned out to be challenging for most groups (with a 95% quantile of GDT_TS for the best model of every group below 60, see **Fig. 3A**), but even in those cases, folds were accurately predicted by the best model (supplemental **Fig. S1**), again suggesting that the folding problem of single proteins is nearly solved.

Similarly to CASP15, there is no single group that stands out^30^. Each group can only generate close-to-best models (GDT_HA difference from the best model < 0.5) in 20% of all EUs (supplemental **Fig. S2**). Many leading groups utilized AF2 and AF3^6,26^ as well as customized MSA inputs to improve the performance of these networks. These customized MSAs are ususally deeper than AF2’s default ones and they are built by searching homologs from additional sequence databases and/or using structure-aided homology searches. In CASP16, no group used pure protein language model (PLM)-based methods, in contrast to CASP15, where two groups used them^30,40^.

A total of 69 groups participating in the monomer structure prediction category submitted an abstract briefly explaining the methods they used in CASP16, allowing us to annotate the strategies used by each group (on top of **Fig. 1**). We partitioned groups based on the types of modeling approaches and compared the performance between different strategies. A majority of groups used AF2 and nearly half used AF3 (**Fig. 2A**). Our analysis indicates that using either AF2 or AF3 yields a better performance than using neither of them, and that only using AF3 or a combination of AF2 and AF3 is better than only using AF2 (**Fig. 2B**). This suggests that AF2/3 remains the state-of-the-art in protein structure modeling and AF3 does show noticeable advantage over AF2. It is worth noting that during the CASP16 prediction period, AF3 was only released as a web server and CASP16 predictors could only control the input sequences (full proteins or segments), test different stoichiometry, and sample more models by registering more accounts (each account can submit 20 jobs per day) and using multiple random seeds. With the recent public release of AF3’s source code and trained weights, we expect further improvement in this field as predictors adopt AF3 in their pipelines and explore better ways of utilizing it. Furthermore, using customized MSAs also correlates with better prediction accuracy (**Fig. 2C**). The observed performance differences are incremental but statistically significant. These findings support that leveraging the power of AF2 and AF3, as well as better MSA, can lead to superior performance in monomer structure prediction.

**Figure 2.**
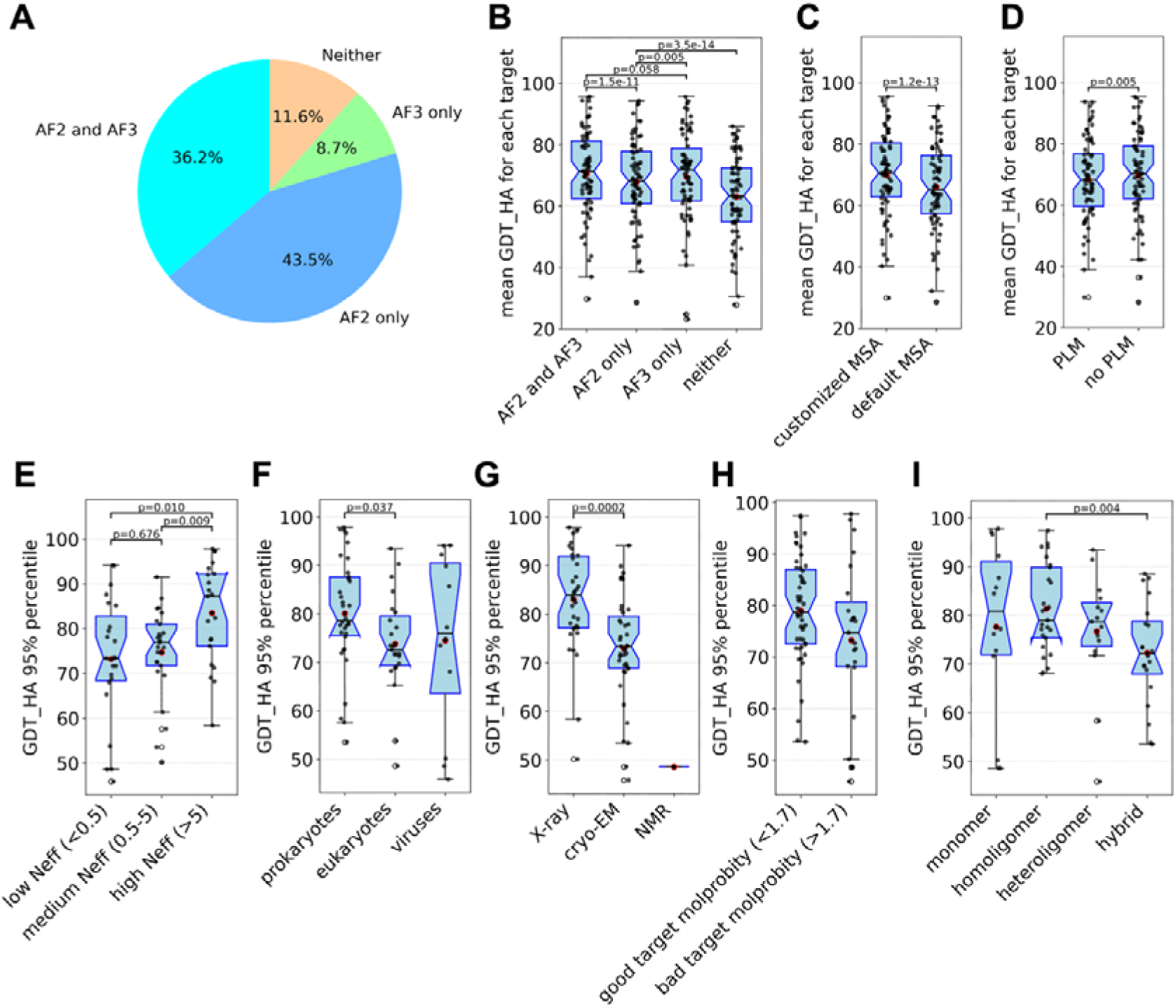
Comparisons of different strategies and correlations between prediction quality and target features. **A)** Of all the groups that provided abstracts and submitted monomer targets, the percentage of AF2 and AF3 usage. **B-D)** y-axis: mean GDT_HA of each target from a subset of groups. x-axis: the partition of groups based on the usage of AF2/3, customized MSA, and PLM. **E-I)** y-axis: 95% quantile of GDT_HA scores for models of a target. X-axis: the partition of targets based on sequence diversity in MSAs, taxonomy, experimental technique, MolProbity quality, and the type of experimental structure it is from.

By contrast, we do not observe a significant difference in performance between groups that used PLMs and those that did not. In fact, groups not using PLMs showed slightly better performance in terms of mean GDT_HA scores (**Fig. 2D**). However, we believe this should not be interpreted as a negative contribution of PLMs; rather, it indicates that better performance can be achieved by optimizing AF modeling strategies (e.g., input sequences, MSAs, broader sampling) without the need for PLMs. Many outstanding groups did not use PLM in CASP16, which aligns with the observation in CASP15 that only using PLM instead of AlphaFold-like networks usually leads to suboptimal performance. In CASP16, PLMs are mostly used to conduct homology search or provide embeddings for AlphaFold-like networks. The optimal role of PLMs in protein structure prediction remains an open and promising area for future exploration.

### EU features affecting performance

We analyzed structural and biological features of EUs that may affect the predictors’ ability to predict structures. We first found that prediction accuracy is positively (but not linearly) correlated with MSA depth measured by *Neff*^41^ (**Fig. 2E**). Deeper MSAs provide rich coevolutionary signals and increase the likelihood of obtaining high-quality models. However, deep MSAs are not always required for high-quality predicted structures. It is observed that AF2/3 can predict 3D structures of a small fraction of natural protein sequences to high quality without MSAs, indicating that AF2/3 also learned additional physicochemical rules about protein folding^42,43^, which might be sufficient to fold a subset of proteins.

Another related factor associated with prediction quality is the phylogenetic origin of a target, which can be broadly categorized into prokaryotic, eukaryotic, and viral proteins (**Fig. 2F**). Generally, prokaryotic proteins yield the deepest MSAs, as a result of broader taxonomy diversity and faster evolution rates compared to eukaryotes^44–47^. Predictors can also effectively mine extensive metagenomic databases to obtain more sequences in addition to sequenced genomes^48,49^. In contrast, eukaryotic and viral proteins tend to have shallower MSAs. Additionally, the rapid evolution of some viral proteins^50^ and their tendency to form obligate oligomers^51^ also make them more challenging targets. Reflecting these expected challenges in generating high-quality models of viral proteins^52^, they were not included in AlphaFold protein structure DataBase (AFDB)^53^.

The experimental method used to determine protein structures also appears to influence prediction accuracy, and the performance on structures solved by X-ray crystallography is generally better. However, the apparent difference between X-ray and cryo-EM could be attributed to the inherent difficulty of the targets—those resolved by cryo-EM are often larger and more complex, which typically leads to lower prediction accuracy. There is only one NMR target in this CASP (T1226-D1, from turkey viral polyprotein), and the performance for that target is particularly poor (**Fig. 2G**). A close inspection of this target reveals that a flexible loop in T1226-D1 (supplemental **Fig. S3B**) is frequently modeled as an ordered alpha-helix. Because structures solved by NMR tend to show higher flexibility compared to those determined by other techniques^54^, we think that the predicted structure might represent a true alternative conformation, a hypothesis that remains to be tested by additional experimental data.

We noticed a couple of targets, such as T1220s1-D1 (supplemental **Fig. S3A**) and T1226-D1 (supplemental **Fig. S3B**) showing low agreement with predicted structures might be poorly resolved and show poor MolProbity scores (>3, and high values indicating more violations to physicochemical rules)^55^. We also found a positive correlation between lower MolProbity scores of targets and poorer agreement with predicted structures (**Fig. 2H**). This is in agreement with another observation that predictors’ performance is better on structures having higher experimental resolution (supplemental **Fig. S3C**). As the performance of structure prediction tools improves, the discrepancy between experimental and predicted structures might be used to detect errors and help to resolve poorly resolved regions in experimental structures^56,57^.

Finally, we found that targets derived from hetero oligomers and hybrid complexes are generally harder to predict than those from monomers and homo oligomers (**Fig. 2I**). Proteins extracted from complex structures frequently undergo conformational changes upon intermolecular interactions and accurate modeling of their structures need to happen in the context of their binding partners. While structure prediction for single-chain proteins is a largely solved problem, 3D structures of many protein complexes remain challenging to predict in the post-AlphaFold era [cite: our oligomer evaluation paper], making monomeric targets derived from these complexes more difficult.

### Potential challenges for current methods

We estimated the field’s performance on each EU by the 95% quantile of GDT_TS among its models, denoted as GDT_TS95. We found that larger EUs tend to show lower GDT_TS95 (**Fig. 3A**). This is primarily because of the nature of GDT_TS calculated on a global scale without normalizing over protein length. Thus, the errors in the 3D space can propagate as the number of residues increases^58^. Taking this EU-size bias into consideration, we identified challenging EUs for most CASP16 predictors as outliers in the GDT_TS95 versus EU size plot (labeled dots in **Fig. 3A**). We manually studied these EUs to identify the weakness of existing methods and suggested future directions to improve structure predictions.

**Figure 3.**
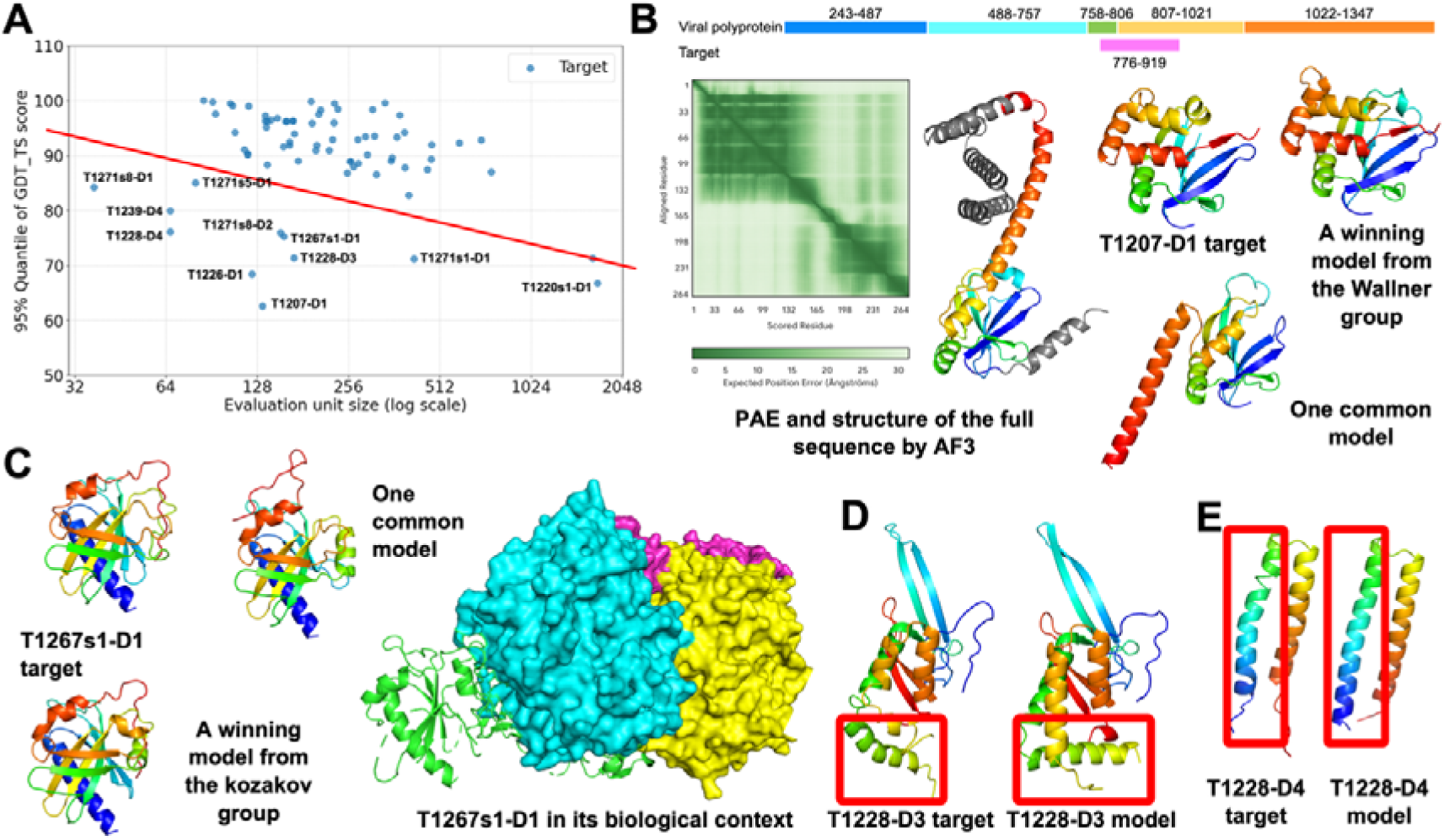
**A)** 95% quantile of GDT_TS scores against target size. The red line is our manual partition of difficulty level. Targets below the line are considered as difficult targets. **B)** Case study for T1207-D1. Top, mapping of the target sequence to its source viral polyprotein. Very few models are correct for this target, possibly due to conformation changes upon sequence truncation. **C)** Case study for T1267s1-D1: predicting its structure requires modeling the whole complex correctly because part of it involves interaction with other subunits. **D)** Case study for T1228-D3: an example of over-stabilizing secondary structures. **E)** Case study for T1228-D4: an example of irregular secondary structures.

#### Potential conformational changes upon truncation

AF2 have been found to be insensitive to mutations and truncations^59,60^. These mutated or truncated sequences frequently do not exist in nature and lack biological relevance, and target T1207-D1 might represent such a case. T1207-D1 originates from an Avian viral polyprotein (**Fig. 3B**). Based on UniProt^61^ annotation, the experimental construct of T1207-D1 spans two proteins (residues 758-806 and 807-1021) cleaved from this polyprotein. Compared to full biological sequences (residues 758-806 and 807-1021), T1207-D1 is a truncated version mapping to residues 776-919 in the polyprotein (**Fig. 3B** top). We predicted 3D structures of the full sequences using the AF3 web server, and the predicted aligned error (PAE) plot and predicted structure suggest that the biological sequences might fold into two domains (**Fig. 3B** left). The T1207-D1 target contains the entire N-terminal domain and part of the C-terminal helical domain. This truncation probably resulted in a conformational change as observed in the experimental structure (**Fig. 3B** middle), where the helical fragment from the C-terminal domain folds back to the N-terminal domain and forms a beta sheet with the N-terminal beta strands. Four out of five models from the Wallner group, as well as a few models from several other groups, resemble the experimental structure, while most other models predicted the segment from the C-terminal domain as an extended helix^29^. We hypothesize that the experimental structure represents a conformational change upon this truncation. A recent study^62^ on similar proteins of T1207-D1 in the Parechovirus family confirmed that there will be a large conformational change upon the truncation of C-terminus residues, supporting our hypothesis. Current methods, being insensitive to truncation, might have failed to predict the conformational changes.

#### Induced fit upon binding

Several most challenging targets from protein complexes contain regions that interact more extensively with other proteins than within the targets. One example is T1267s1-D1 (**Fig. 3C**), a component of the SPARDA complex, which plays an important role in the bacteria defense system. While most groups were able to predict the structured regions of this target, they failed to accurately predict the orientation of the loop mediating its interaction with a binding partner. This loop might undergo induced fit upon binding, and its conformation can only be correctly predicted in the context of its parent oligomeric target H1267, which is a challenging target only correctly predicted by a couple of groups, including the kozakovvajda group. In addition, EUs T1271s1-D1, T1271s5-D1, T1271s8-D1, and T1271s8-D2 are derived from a large RNA editing core complex, and they suffer from similar issues (see the CASP16 website for additional details). To investigate whether interface residues are generally harder to model, we computed the GDC_mc scores for interface and non-interface residues for all models that have overall GDT_TS above 80. Residues at interfaces between chains or EUs display lower GDT_mc scores compared to non-interface residues (supplemental **Fig. S4**).

#### Irregularities in secondary structures

Consistent with previous observations^63,64^, we noticed a tendency of over-stabilizing helices or strands in flexible regions, and incorrectly predicting irregular secondary structures such as bent helices, smaller helices (3□□ helix), and larger helical turns (π-helix). Two poorly predicted small EUs, T1228-D3 and T1228-D4, from serine-type integrase SprA contain slightly bent □-helices. Most predictors missed such irregularities and predicted longer and straight helices instead (**Fig. 3D** and **3E**). This is possibly related to the fact that the training data of AF2/3 are biased towards ordered structures and regular secondary structures. We analyzed all the targets and top 10% models of each target with MolProbity. We found that compared to models, target structures tend to have more Ramachandran outliers than these high-quality models (supplemental **Table S2**), supporting our observation that irregular secondary structures in target structures are hard to capture in predicted structures. Additionally, these irregularities in secondary structures in a target might not be conserved in its homologs (for example, a homologous target, T1239, does not have these irregularities). Therefore, coevolutionary information derived from MSAs might be misleading as well.

### Performance evaluation and ranking

#### Scoring formula

We investigated a number of classic prediction assessment measures. To understand the relationships among various measures, we conducted correlation analysis and factor analysis (supplemental **Fig. S5**). These analyses allowed us to group the evaluation metrics into categories, from which nine representative measures were selected: **GDT_HA, QSE, reLLG_const, SphGr, CAD_AA, GDC_SC, AL0_P, lDDT,** and **MolProbity score**. In Methods, we briefly describe each score.

Adhering to the CASP protocol, we computed z-scores (see Methods) for each selected measure on models of each EU. To avoid over-penalizing failed or missing predictions, we imputed z-scores below -2 and missing values as -2. An alternative strategy frequently used in CASP is to set z-scores below 0 to 0. We tried both strategies in our assessment and prioritized the former because it gives us better discrimination between groups. In an era when most targets are correctly predicted and the difference in performance between groups is subtle, being able to consistently generate high-quality structure for every target is important.

The total z-score of a prediction is computed as the weighted sum of z-scores of the nine measures. The assigned weights to each measure reflect both precedents from previous CASP experiments and our judgment regarding the importance of each measure. For example, the weight of GDT_HA, QSE, and reLLG_const being 1/6 is consistent with CASP15^30^. The higher weights on these scores emphasize three aspects: the importance of overall structure^58^, quality evaluation, and how these models can help experimentalists in molecular replacement^65,66^, respectively. The other 6 measures are given either 1/8 or 1/16 weight, depending on their relative importance in estimating model quality.

It is important to note that due to the high correlation between different measures (supplemental **Fig. S5** top), the ranking, especially the selection of the winning group, is largely not sensitive to the change of weights. For example, changing the weight of a component score to 0 will result in a very similar ranking, and the top groups will largely remain the same (supplemental **Table S3**). Only changing the weight of the Molprobity score to a higher value will significantly change the ranking because Molprobity scores do not correlate with structural similarity between a target and a model, and different groups vary greatly in MolProbity scores of their models. While it is meaningful to use Molprobity scores to encourage people to mind the geometric accuracy of predicted structures, we chose to use a low weight, as in CASP15, to prevent it from dominating the ranks.

Each monomer target in CASP16 contains one to four EUs, and two targets with the largest number of EUs (T1228 and T1239) share similar 3D structures. To prevent targets with more EUs from dominating the evaluation, we assign a weight to each EU based on the number of EUs per target (i.e., a weight of 1/N for each EU if one target has N EUs), resulting in an equal weight for each target. The final scoring formula is presented below.

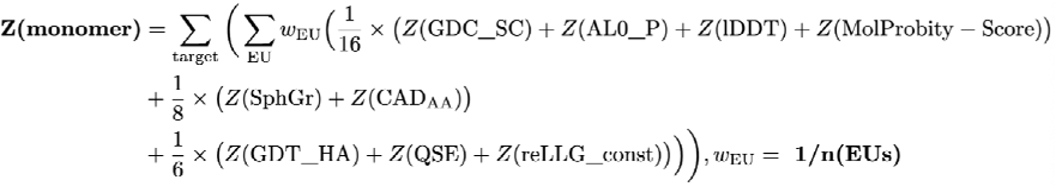

#### Phase 1 ranking

We consider Phase 1 the primary phase of CASP16 due to the following reasons. First, its setup is the most similar to traditional CASP experiments; second, this phase contains the largest number of EUs; third, predictors show the highest participation rate in this phase. We ranked groups by the aforementioned formula over all Phase 1 EUs. Additionally, we conducted a head-to-head comparison with bootstrapping (see Methods) to provide statistical significance for performance differences between groups. The leading groups in Phase 1 are Yang, Yang-Multimer, Yang-Server, MULTICOM, and MULTICOM_human [cite: CASP16 paper from Jianyi Yang and Jianlin Cheng], which significantly outperform the rest (**Fig. 4A** and **4B**). By looking at the contribution of individual scores, we noted that the winners typically excel in measures like GDT_HA and reLLG_const. However, they often score lower in MolProbity, suggesting that their models may violate more physicochemical constraints. Notably, although these models that ranked the top are relatively poor in MolProbity scores among all models, the MolProbity scores of experimental structures are not better top 10% models for each target (supplemental **Table S2**). Nevertheless, we encourage future participants to pay greater attention to the physical validity of their models. Two of the top-performing groups in CASP15, PEZYFoldings and UM-TBM (from Zheng lab), are less competitive in CASP16.

**Figure 4.**
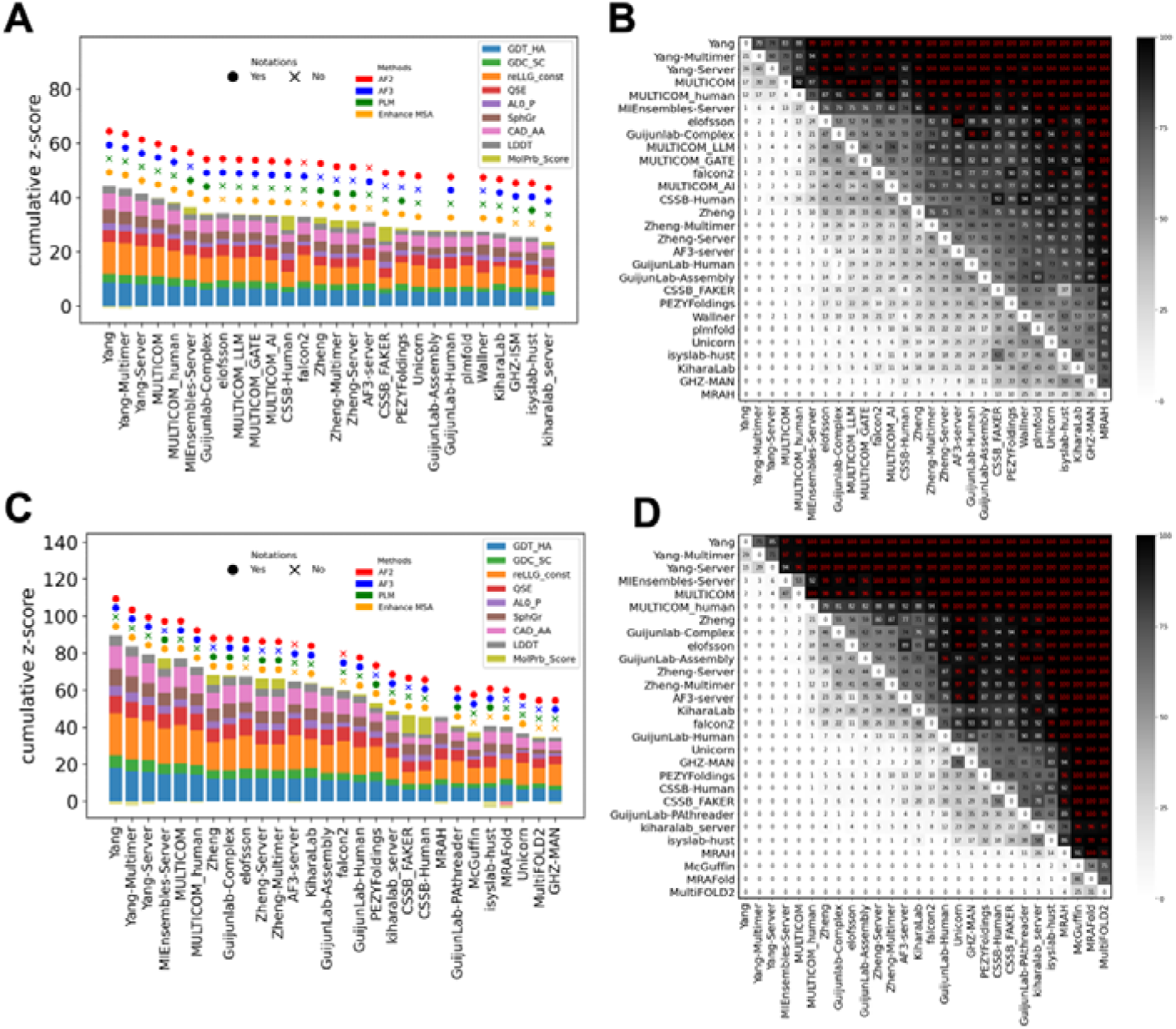
**A)** Ranking using all Phase 1 targets and best models. **B)** Results of head-to-head comparisons with 1,000 bootstrapping samples. The values in each box reflect the percentage of bootstrap samples where a group (labeled on the Y-axis) outperforms another one (labeled on the X-axis). Red font in each black cell indicates significance at a confidence level of 0.05. Bootstrapping is done only between the groups present in **A** and is ranked based on the row sum. **C)** Ranking using all Phase 0, 1, 2 targets and best models. **D)** Similar to B, but done on targets from all phases and using groups present in **C**.

#### Main ranking based on targets from all phases

CASP16 introduced two new Phases— Phase 0 and Phase 2—which expanded the scope of monomer structure prediction. We combined targets from all three phases to determine the overall ranking for CASP16 monomer prediction and considered this as the main ranking. All three groups from the Yang lab [cite: CASP16 paper from Jianyi Yang] had the highest z-score, which aligns with the results from Phase 1, and their superior performance is statistically significant (**Fig. 4C** and **4D**). This consistency highlights the robustness of their superior performance across prediction Phases in CASP16, and we consider the Yang lab as the winner for monomeric structure prediction. A unique strategy, heavily used by the Yang lab is to refine the input sequences by removing the potential disordered regions and using partial (fragments) instead of the full sequences as inputs of AF2/3. We termed this strategy fragment-based modeling, which resembles the process of experimental construct refinement widely used in experimental structure determination. Fragment-based modeling seems to be a successful strategy in improving modeling quality.

#### Phase 0 ranking

Phase 0 is designed to challenge predictors’ ability to model oligomeric complexes by not providing stoichiometry information to them. Incorrectly predicted stoichiometry and complex structures might affect the prediction accuracy of monomers derived from these complexes, because interaction with other partners might affect the conformations of these monomers. Phase 0 rankings differ noticeably from those in Phase 1, but surprisingly, groups that performed the best in Phase 0 monomer targets (supplemental **Fig. S6A**) are not the winners of Phase 0 in the oligomer category. Instead, several groups that did not utilize AF3 (NKRNA-s, MIEnsembles-Server, Zheng-Multimer, and Zheng) were ranked at the top for Phase 0 monomer targets. We speculate that this is because the winners (e.g., the Yang lab) of the monomer category relied on extensive use of AF3 to obtain better models, but they were limited by the quota per user implemented in the AF3 web server. Because some Phase 1 targets’ sequences were released during Phase 0, and all Phase 2 targets’ sequences were released during Phase 1, they had more time to gather AF3 models for the two phases coming after Phase 0, where they outperformed other teams. They might also have chosen to prioritize Phase 1 targets in modeling via the AF3 web server.

#### Phase 2 ranking

Phase 2 is designed to evaluate the predictors’ ability to utilize the large number of 3D models generated by MassiveFold in their prediction pipeline. However, the integrity of the Phase 2 experiment may have been compromised, as the winning teams likely conducted extensive sampling using their own pipelines in combination with the AF3 web server. The groups from Yang Lab demonstrated an even greater advantage in Phase 2 compared to earlier phases (supplemental **Fig. S6B**), consistent with the observations in the following sections that the groups from Yang Lab are still able to improve in Phase 2. This result may reflect their superior ability to effectively leverage large amounts of AF2/3 models to improve prediction quality or a more extensive usage of AF3 to iteratively refine the input sequences for better model quality [cite: CASP16 paper from Jianyi Yang].

#### Rankings partitioned by difficulty

In the earliest CASP exercises, there were at least two categories in structure prediction: comparative modeling and *ab initio* (new fold)^1,67,68^. These two categories further became what is known today as template-based modeling and template-free modeling^69,70^. Before the use of coevolution-based contact prediction and the emergence of deep learning methods^71^, having homologous structures in the PDB was critical for correctly predicting the fold of a target^72,73^. However, as modern deep learning methods have revolutionized protein structure prediction techniques, having templates detectable by homology search is no longer necessary for obtaining high-quality models. Notably, AF2’s performance on template-free targets approached that of easier ones^9^, making the traditional dual-category assessment unnecessary. Consequently, since CASP15, the organizers merged these categories into a unified evaluation of single protein modeling^30^.

We maintain this unified approach in CASP16. We noticed that there was no significant correlation between the availability of PDB templates (found by HHsuite^74^) and model quality (supplemental **Fig. S7**). However, it remains meaningful to emphasize groups’ performance on targets that are challenging. As shown in **Fig. 3A**, targets remaining difficult to the field can be separated from the rest based on GDT_TS95, and we considered targets below the red line in **Fig. 3A** as difficult and the rest as easy. We evaluated groups’ performance on the difficult and easy targets separately and the results are shown in supplemental **Fig. S8**. Notably, groups such as Wallner, CSSB-Human, and Yang performed relatively well on difficult targets. In particular, both the Wallner and the CSSB-Human groups were able to correctly predict the topology for the aforementioned challenging target, T1207-D1 (**Fig. 3B**), significantly outperforming other groups on this target. Additionally, CSSB-Human achieved a relatively better performance in MolProbity score, which also contributes to its top ranking in the category of difficult targets.

#### Sidechain accuracy evaluation

Our initial assessment of sidechain accuracy employed the GDC_sc scores provided by the prediction center. However, the Phase 1 target ranking derived from GDC_sc z-scores (Supplemental **Fig. S9** top) exhibited a strong resemblance to the overall model quality ranking (**Fig. 4A**), with the Yang lab groups consistently performing best. We were concerned that this high correlation might be attributed to GDC_sc being a superimposition-based metric, inherently making it dependent on backbone accuracy and therefore strongly correlated with other superimposition-based scores such as GDT_TS30.

To address this, we implemented an alternative sidechain quality evaluation method that operates independently of backbone modeling quality. Following a methodology similar to CASP15’s evaluation, we employed the Average Absolute Accuracy (AAA) score, initially detailed in the SCWRL4 paper. The AAA score is calculated for each residue, indicating the percentage of the model’s χ-angles that deviate by no more than 40° from their counterparts in the target structure. A model’s comprehensive AAA score is then computed as the average of these per-residue scores. Critically, because the AAA score relies exclusively on χ-angles, it remains independent of backbone conformation. Despite this independence, the top-performing groups in AAA scores largely remain the same as those in the main ranking, though the absolute differences between groups are subtle (Supplemental **Fig. S9** bottom). This outcome suggests that the accurate prediction of sidechain rotamers necessitates precise prediction of their surrounding environment, thereby implying a fundamental requirement for accurate backbone modeling.

### Insights from additional CASP16 experiments

#### Comparisons between different phases

In CASP16, each successive phase was designed to provide predictors with additional information with the aim of reducing the difficulty and stimulating better models. We wondered if CASP16 groups could successfully utilize such additional information, and thus we compared each group’s performance difference between consecutive phases (in GDT_HA), focusing on shared targets between them. We focused on 28 groups showing superior performance in our final ranking (targets across 3 phases). For the comparison between Phase 0 and Phase 1, we focused on the first models because the first models reflect each group’s best estimate of stoichiometry. It is common for the predictors to use a diverse range of predicted stoichiometry in all five models, and it will be less informative to use the best models here.

Overall, providing the stoichiometry information in Phase 1 and providing over 8,000 MassiveFold models in Phase 2 helped over 60% groups to improve their performance (**Fig. 5A** and **5B**), but such improvement is more noticeable for Phase 1. The smaller improvement in Phase 2 is likely related to two additional factors. First, many groups possibly emphasized more on Phase 1 than the other two phases. Second, the winning groups mostly have their own pipelines to perform extensive sampling through AF2 and AF3, thus, providing additional MassiveFold models is not necessarily helpful; instead, people had more time to sample more AF3 models in Phase 2, which could have been more important. Below, we highlight several cases that are indicative of the differences between phases.

**Figure 5.**
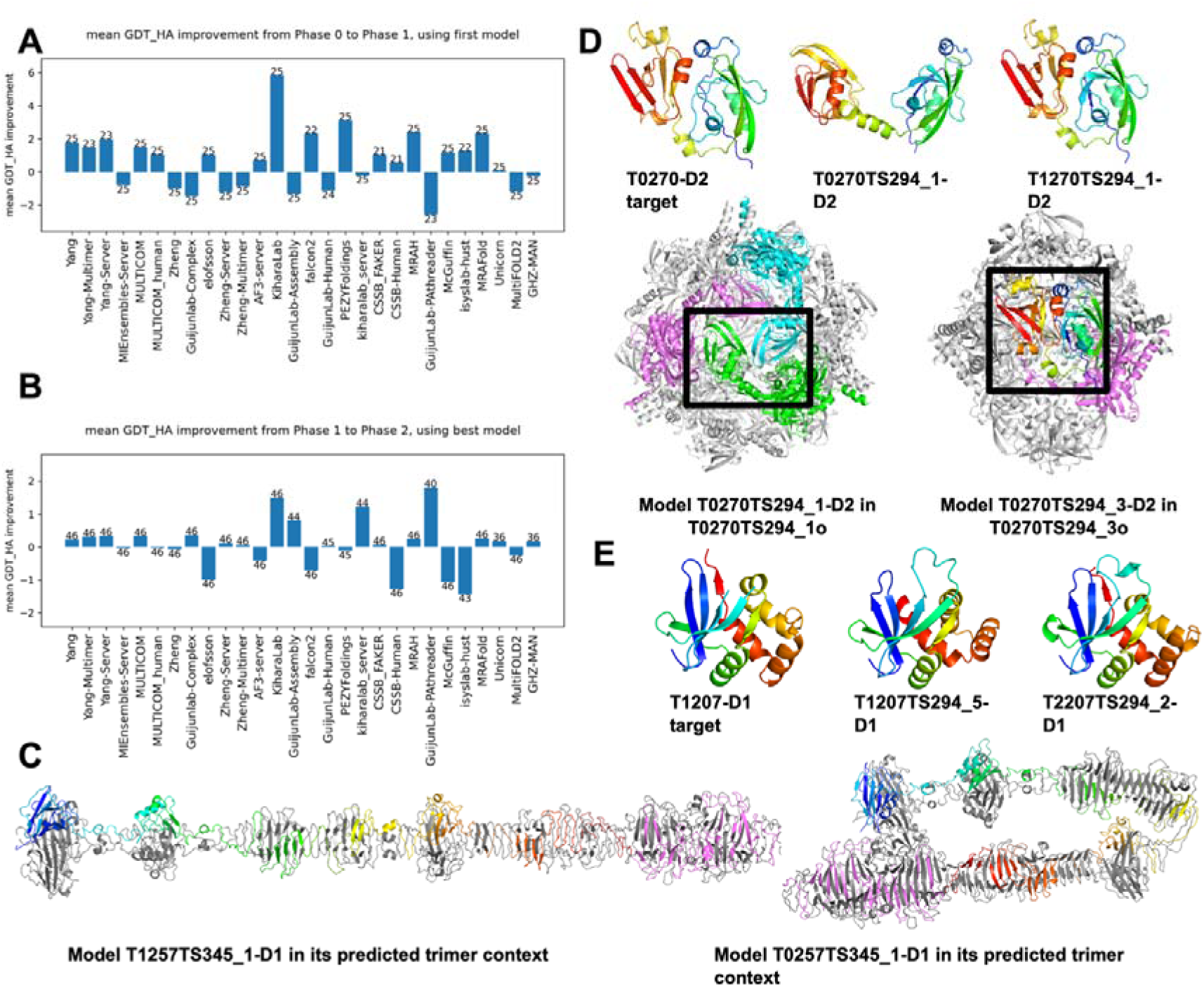
**A)** Average change of GDT_HA for all the targets passed from Phase 0 to Phase 1. The number on each bar indicates the number of targets present in both Phases submitted by this group. **B)** Average change of GDT_HA for all the targets passed from Phase 1 to Phase 2. The number on each bar indicates the number of targets present in both Phases submitted by this group. **C)** Left: a correct model in Phase 1; right: an incorrect model that bends this long protein. **D)** Top: target and model structure for T0270-D2 and T1270-D2; bottom: the predicted models in their oligomeric context. **E)** Target and model structure for T0270-D2 and T1270-D2.

#### Heavier human involvement improves structure prediction quality

because all Phase 0 targets were reused in Phase 1 and predictors could invest more time and apply human intervention to improve prediction quality. An example of large improvement comes from the models for targets T0257-D1 and T1257-D1 from the MULTICOM group (**Fig. 5C**). T0257-D1/T1257-D1 is from a trimer of the side tail fiber protein pb1 of bacteriophage. Predicting the stoichiometry of such complexes should be trivial based on AF2/3 prediction and homology-based inference. However, a significant improvement is observed due to the selection of a straight model (**Fig. 5C** left) over a bent one (**Fig. 5C** right) by the MULTICOM team in Phase 1. Both AF3 server and ColabFold have a tendency to bend such elongated proteins and are not likely to generate the correct conformation automatically (supplemental **Fig. S10**). However, knowledge about viral tail proteins indicates that they should typically adopt highly straight and elongated conformations. Thus, in Phase 1, the MULTICOM team modeled the protein as 3 segments (with overlaps), and these 3 segments are all predicted to have a straight conformation. They then superimpose the overlapping regions to obtain a straight conformation over the entire protein [cite: CASP16 paper from Jianlin Cheng].

#### Knowing stoichiometry helps people to select the correct monomer conformation

Another example of extreme improvement in Phase 1 is between target T0270-D2 and T1270-D2 (**Fig. 5D**). These targets are of HtrA serine protease from *Borreliella burgdorferi*, a pathogen causing Lyme disease. In Phase 0, the KiharaLab group [cite: CASP16 paper from Daisuke Kihara] sampled multiple stoichiometry and selected a 12-mer (**Fig. 5D** bottom left) as their first model. However, this target should be a hexamer instead. In Phase 1, the same group selected a hexamer as model_1 (**Fig. 5D** bottom right). Although this hexameric model is also suboptimal in its oligomeric interfaces, it allowed them to correctly predict the monomer conformation. Comparison between the incorrect Phase 0 prediction (**Fig. 5D** top middle) and the correct Phase 1 prediction (**Fig. 5D** top right) revealed that the former involves domain swapping between interacting chains, resulting in a less compact monomer model.

#### Extensive sampling improves model quality for a challenging target

The challenging target T1207-D1 (**Fig. 3B**) demonstrates the benefit of extensive sampling. In Phase 1, only very few groups, especially the Wallner group which performs extensive sampling using AFsample^29,75^, were able to correctly predict its structure. However, leveraging the extensive models provided in Phase 2, other groups, for example, KiharaLab, were able to identify the correct conformation (**Fig. 5E**). Using large structure pools and combining several scores based on statistical potentials, KiharaLab were able to improve their Phase 2 predictions compared to Phase 1 [cite: CASP16 paper from Daisuke Kihara]. These observations suggest that extensive sampling, either by MassiveFold or by the predictors themselves, in combination with suitable scoring functions, can enhance structure prediction accuracy.

#### Evaluation of Model 6

Model 6 predictions are restricted to use MSAs generated by ColabFold, allowing us to separate the contribution by improved MSAs and advanced structure prediction deep learning networks. To assess both the quality of MSAs and the network architectures, we conducted two comparative analyses: 1) model 6 vs. model 1 from the same group – to evaluate the performance of each group’s own MSA construction strategy; 2) model 6 vs. ColabFold model – to assess how the group’s modeling network performs relative to ColabFold, given the same MSA input. Groups such as Zheng, MULTICOM_human, MULTIFOLD, and PEZYFoldings [cite: CASP16 paper from Zheng, Jianlin Cheng’s and PEZYFolding’s team] are able to obtain better results for model 1 than model 6, indicating that their group’s MSA is better for structure predictions (supplemental **Fig. S11** top). Meanwhile, groups such as Guijunlab-QA, Guijunlab-Human, and MULTICOM [cite: CASP16 paper Guijun Zhang and Jianlin Cheng], were able to outperform ColabFold using the same MSAs, although such an advantage appears to mainly come from several targets that ColabFold struggled with (supplemental **Fig. S11** bottom).

Although these 6th models were designed to provide invaluable insights about strategies used in the field, they were not considered in the main CASP16 rankings, and consequently, few groups submitted model 6 predictions. Even among those groups that did, many model 6 submissions were missing for a large number of targets. As a result, the available data for model 6 is too limited to support any robust analysis or strong conclusions. We strongly encourage future CASP participants to submit more comprehensive model 6 predictions. By doing so, it would allow for a more systematic and quantitative evaluation of how MSA quality and innovations in model architectures independently contribute to structure prediction performance.

### Quality evaluation

As the field advances in modeling monomer structures and large-scale model generation, the ability to rank predicted structures and predict model quality becomes more important in this post-AlphaFold era. We thus performed a series of analyses aiming at evaluating CASP16 groups’ ability in self-assessment and model selection.

#### Ranking by the first model

The first model represents the structure that the predictor is most confident in and thus can serve as an indicator of both prediction quality and model selection ability. Interestingly, if we evaluate the performance of CASP16 groups by the first models, the Yang lab maintains the top position, but the AF3 server rose to second place (**Fig. 6A**), showing a performance not statistically worse than the top group (**Fig. 6B**). Yang’s high rank can likely be attributed to its superior modeling accuracy, whereas AF3’s increase in rank is likely to be driven by its improvement in model evaluation and ranking capacity.

**Figure 6.**
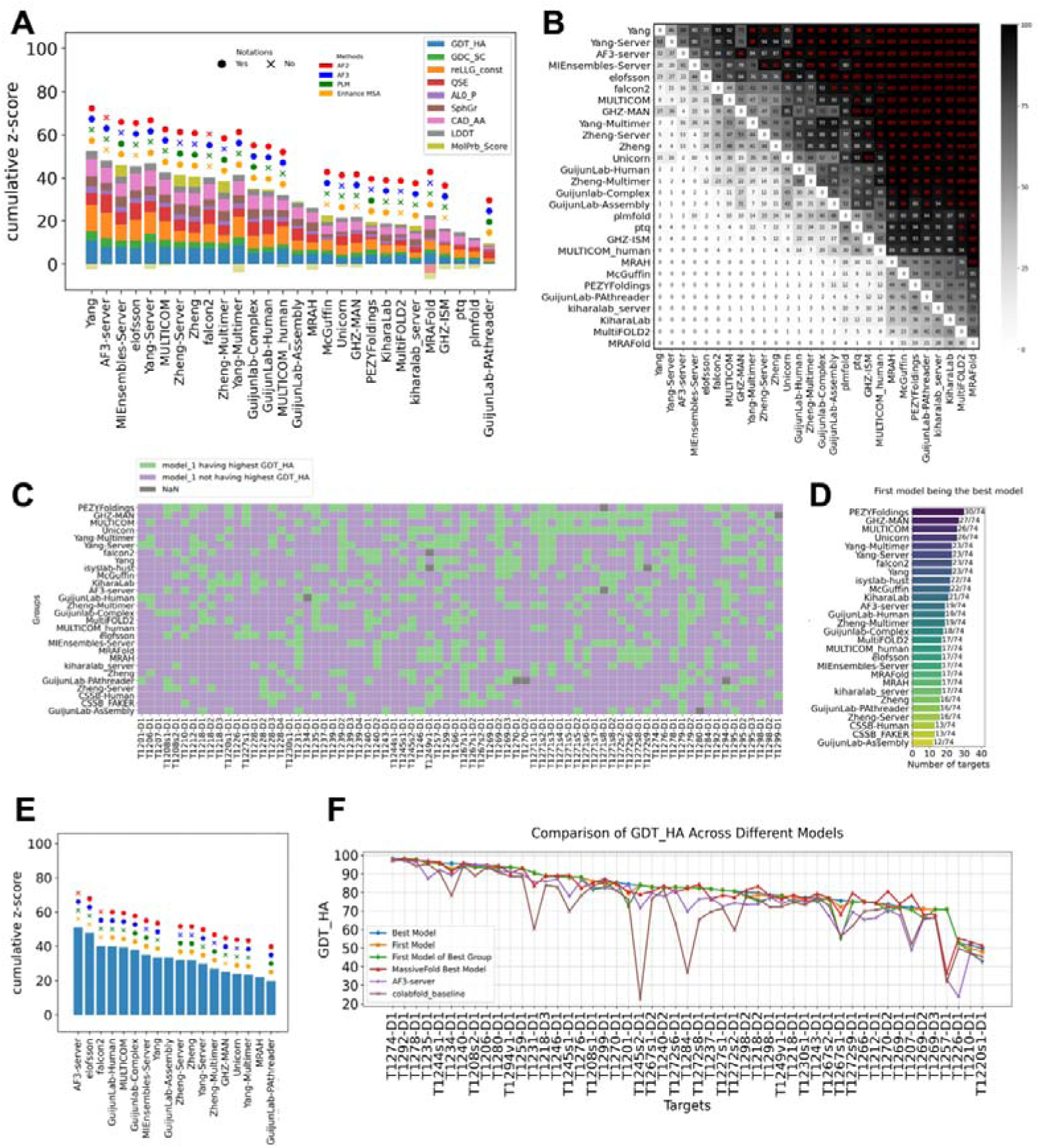
**A)** Ranking using Phase 1 targets and first models. **B)** Results of head-to-head comparisons with 1,000 bootstrapping samples. The values in each box reflect the percentage of bootstrap samples where a group (labeled on the Y-axis) outperforms another one (labeled on the X-axis). Red font in each black cell indicates significance at a confidence level of 0.05. Bootstrapping is done only between the groups present in **A** and is ranked based on the row sum. **C)** Statistics on the frequency of the first model having the highest GDT_HA across all models for every target. Only the top 28 groups in the main ranking are presented. **D)** Ranking based on the count of green cells (targets where the first model is the best one for a group) in each row of **C**. **E)** Ranking using only QSE scores. **F)** Values of the highest GDT_HA among MassiveFold models or different sets of models submitted by CASP16 groups. Targets are ordered by the highest GDT_HA from left to right.

#### Frequency of first models being the best models

To directly evaluate each group’s ability in selecting the first model, we calculated the percentage of cases in which a group’s first model was also the best among the five submitted models for that target. Among the top-ranking groups, many can do only slightly better than a random selection, where the first model has a 20% chance of being the best (**Fig. 6C** and **6D**). One group, PEZYFoldings, was able to reach a ratio of 30/74 (**Fig. 6D**), highlighting a robust ability in ranking its own models effectively. This is also consistent with their best performance in selecting the first model for oligomeric targets [cite: our oligomer evaluation paper].

#### Ranking by QSE (a.k.a. ASE) scores

Starting from CASP15, predictors are required to submit their own “pLDDT score” in the b-factor column of their predicted structures^33^. QSE score is subsequently developed to measure the agreement between pLDDT and the real LDDT based on the comparison between a model and a target. By ranking only using QSE scores, we discovered that the AF3 server obtained the highest QSE scores, followed by a couple of other groups, i.e., elofsson and falcon2, that explicitly rely on the AF3 server (**Fig. 6E**). Together with AF3 server’s near-top performance in the first model quality, our results suggest the complicated confidence module developed in AF3 does remarkably improve its ability in confidence estimation.

#### Analysis of MassiveFold results

One key challenge in structure prediction is finding the best models among the diverse structures sampled by modeling with different input MSAs and structure prediction networks. Utilizing the large number of predicted structures by MassiveFold for most CASP16 targets (monomers extracted from protein-nucleotide hybrid complexes did not have MassiveFold results), we were interested in the following question: will we obtain a better prediction among the MassiveFold models if we have a perfect selector to rank them? To address this, we ran LGA for all models from the MassiveFold against their corresponding target. We plotted the best GDT_HA for MassiveFold models on each target against the best models among the around 300 predictions submitted by CASP16 groups for each target.

The best models generated by CASP16 groups are generally better than the best MassiveFold models, suggesting the innovations in modeling strategies used by CASP16 groups resulted in a meaningful improvement in model quality. The most extreme case of human improvement happened to target T1257-D1 discussed above (**Fig. 5C**), a target where automatic predictions from MassiveFold tend to bend the elongated structure. We also noticed that MassiveFold models tend to have worse performance for large targets. Two reasons are likely behind this observation: first, the large complexes are likely harder for automatic prediction without human guidance; second, MassiveFold also generated fewer models (2,040 or 4,040, instead of 8,040) for these large targets due to limits in computational resources.

However, the pool of structures generated by MassiveFold did contain better models for some targets, indicating that our ability to model monomer structures can be further improved by optimizing target selection metrics. Notably, MassiveFold predictions were generated based on AF2-like networks. The diffusion module used by AF3 made it intrinsically suitable for generating a diverse set of predictions and enlarging the pool of possible models to select from. Thus, we expect that further performance improvement can be enabled by better quality assessment and model ranking methods, highlighting self-evaluation as a critical direction for future improvement.

### Lack of obvious progress in monomer prediction

To evaluate whether the field has made progress from CASP15 to CASP16, we compared the predictors’ performance in CASP16 against CASP15. Because the apparent performance of different CASP experiments will be significantly affected by the overall difficulty of targets, we need a method that has not changed between the two CASP experiments to serve as an indicator of target difficulty. We used ColabFold’s performance as an indicator of target difficulty. ColabFold pipeline did not change between CASP15 and CASP16 except updating sequence/structure databases and versions of AF2 and AFM weights (from 2.0 to 2.3). ColabFold submitted predictions to all 112 CASP15 monomer EUs, and 59 out of 74 CASP16 monomer EUs in Phase 1: EUs derived from hybrid targets (with nucleotides) cannot be modeled by ColabFold and were excluded. Accordingly, we restricted our comparative analysis to EUs with ColabFold predictions.

We computed the average GDT_HA scores of the best models from the top 50 groups in CASP15 and CASP16 (**Fig. 7A**). The highest GDT_HA among CASP participants increased from 71.6 to 77.4. However, the GDT_HA of ColabFold also increased—from 64.7 in CASP15 to 72.2 in CASP16—suggesting a potential shift in the overall difficulty of the targets. The distribution of GDT_HA scores across EUs further supports this interpretation, with CASP15 containing a higher proportion of difficult EUs compared to CASP16 (**Fig. 7B**). The difference in target difficulty between the two CASP experiments prevents direct comparison of performance, calling for more robust statistical analysis that can take target difficulty levels into account. One obvious difference is a greater number and fraction of groups outperformed ColabFold in CASP16, indicating the field is surpassing AF2, possibly due to the contribution of AF3.

**Figure 7.**
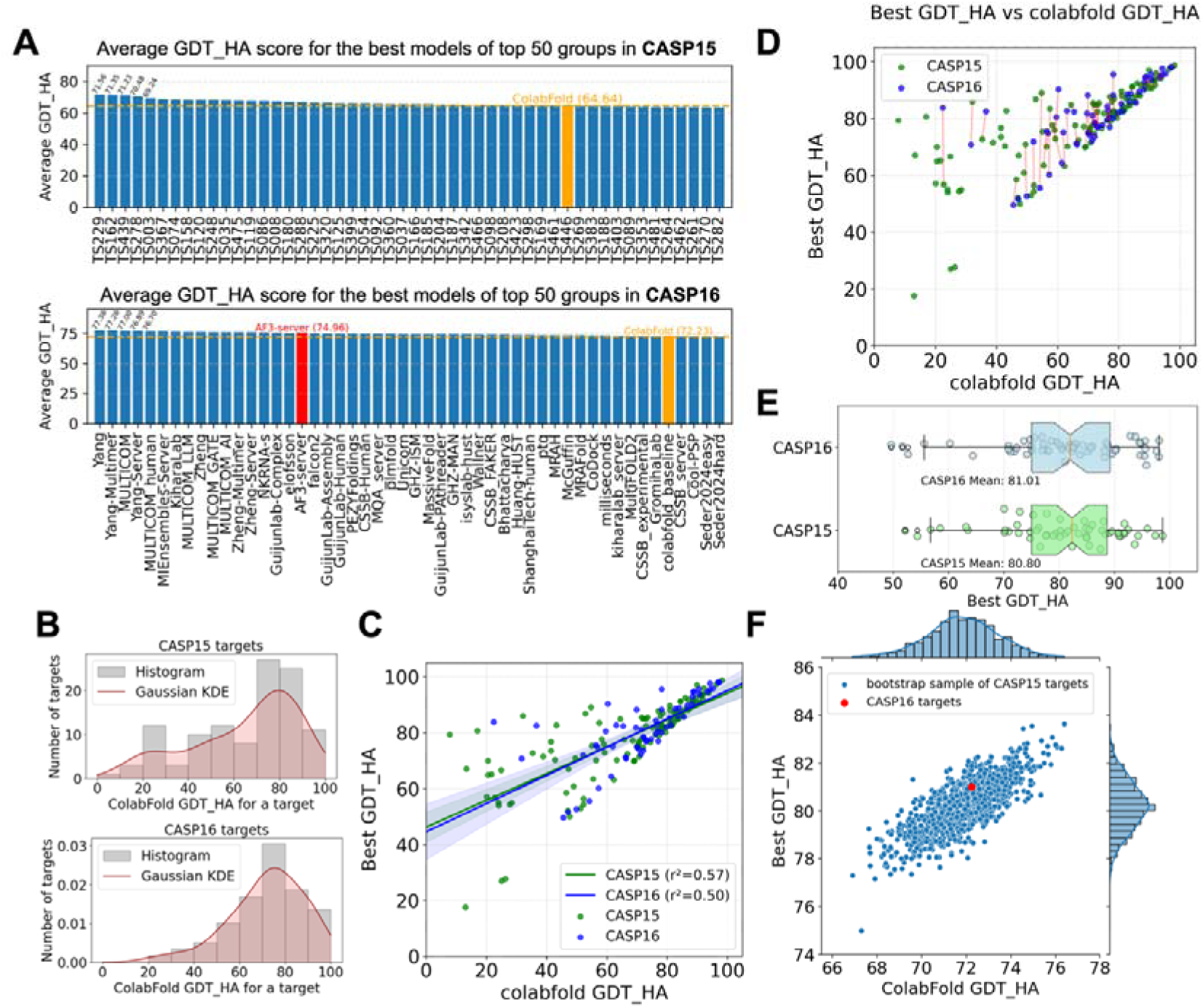
**A)** Average GDT_HA for different groups over all targets submitted by colabfold_baseline for CASP15 (top) and CASP16 (bottom). **B)** Distribution of CASP15 (top) and CASP16 (bottom) colabfold_baseline’s best GDT_HA. **C)** Best GDT_HA among CASP15 and CASP16 groups and colabfold_baseline’s best GDT_HA. **D)** Mapping CASP15 models to CASP16 models to minimize the difference in colabfold_baseline’s GDT_HAs (target difficult measure) between the two CASP experiments. **E)** Best GDT_HA for the mapped targets between CASP15 and CASP16 based on the 1-on-1 mapping in **D**. **F)** Performance comparisons between CASP16 targets and weighted bootstrap samples of CASP15 targets. These bootstrap samples of CASP15 targets show similar difficulty levels (by colabfold_baseline GDT_HA, x-axis) and best performance (by best GDT_HA, y-axis) as CASP16 targets.

To evaluate the progress in a more robust fashion, we first studied the relationship between target difficulty, as indicated by the GDT_HA score of ColabFold’s first model, and the field’s optimal performance (in CASP15 or CASP16) on a target (**Fig. 7C**). Differences in target difficulty can confound evaluation of the progress. Easier targets offer limited room for improvement over the ColabFold baseline, whereas harder targets correlate with bigger performance differences between the best model and the ColabFold model (a potential improvement). To more objectively assess progress between CASPs, we implemented two statistical approaches to mitigate biases arising from variation in target difficulty.

We applied the Hungarian algorithm^76^, which finds the optimal one-to-one matching between two sets, to select a subset of CASP15 EUs to match the difficult level of CASP16 EUs as much as possible. In our case, the set of matched EUs (connected by the red lines in **Fig. 7D**) shows very similar overall difficulty levels between CASP15 and CASP16, with average ColabFold GDT_HA of 72.06 and 72.23, respectively. Based on this matching to eliminate the differences in target difficulty, we calculated the best performance of other groups in CASP15 and CASP16 (**Fig. 7E**) and discovered that the performance in CASP15 and CASP16 is very similar (mean best GDT_HA being 80.80 and 81.01, respectively) without any statistical significance in performance difference (p-value = 0.92).

As an alternative approach, we used a weighted bootstrap technique to generate samples (with replacement) of CASP15 EUs while trying to make each bootstrap sample have a similar ColabFold GDT_HA distribution as CASP16 EUs. For each bootstrap sample of CASP15 EUs, we computed their average ColabFold GDT_HA (target difficulty) and the average GDT_HA of the best model of each EU, and the distribution of these two measurements are shown in **Fig. 7F**. Among CASP15 bootstrap samples with similar overall difficulty levels (near the red dot representing CASP16), about 1/3 show better overall performance than CASP16. This alternative analysis again indicates that although the best performance in CASP16 is slightly better than CASP15, this improvement is subtle and not statistically significant.

Given AF3 allows people to include small molecules in their models of proteins, we wondered whether using AF3 allowed CASP16 groups to predict atomic details around the functional sites (such as metal and small molecule binding sites) to higher accuracy. We thus focused on residues around metals and biological ligands found in structures submitted to the prediction center by experimental labs. We evaluated the prediction accuracy of sidechains of these ligand-binding residues using GDC_all^58^ and compared the performance between CASP15 and CASP16 (supplemental **Fig. S12** and **S13**). Again, we did not notice any remarkable improvement in the modeling accuracy for these important functional sites.

### Inter-domain interaction evaluation

Inter-EU interaction prediction was conducted by monomer or assembly assessors in the previous CASPs^7,77^. Before the recent breakthrough in protein structure modeling, the community struggled in predicting domain-domain interactions (DDIs) without homologous PDB templates. Thus, an EU in early CASPs frequently corresponds to a domain. As the community’s ability to model protein structures continues to grow^7,77–80^, the accuracy in modeling DDIs has drastically increased. In recent years, multiple interacting domains have been frequently merged into a single EU, representing a unit whose structure can be predicted as well as single domains by the community^39^. With the improvement in modeling inter-domain interactions, the size of EUs in CASP also increased.

Among CASP16 monomer targets, we identified 63-68 pairs of DDIs based on Chainsaw (for domain parsing) and PISA (for interface analysis). The uncertainty in the number of DDIs is due to some proteins (T1228 and T1239) adopting multiple conformations and showing variability in inter-domain interactions. 50 (74%–79 %) of these interacting domain pairs were included in the same EU, suggesting that the majority of DDIs can be accurately modeled by the field. The remaining 13-18 DDIs were partitioned into different EUs, and it is not clear whether their relative orientations are intrinsically flexible or they represent challenging cases for DDI modeling.

To probe whether these inter-EU DDIs are of biological relevance, we used PISA^81^, a well-established software that judges whether an observed interface in an experimental structure is biologically relevant, to assess the interfaces between these inter-EU DDIs. As a control, we also used PISA to evaluate the interfaces between DDIs within the same EU. Plotting the interface sizes versus PISA P-value (a measure of biological relevance of an interface), we found that inter-EU DDIs tend to have smaller interface sizes and higher PISA P-value. A PISA P-value of greater than 0.5 is suggested to be indicative of experimental artifacts instead of biologically relevant interactions. Only 5 inter-EU DDIs show a PISA P-value < 0.5, suggesting that they could be biologically relevant and suitable targets for evaluating DDIs. However, manual inspection suggests only three of them are not suitable targets for DDI prediction due to different reasons.

In particular, two EUs, T1239-D2 and T1239-D4 (orange circle in **Fig. 8A** and **Fig. 8B**), might interact or not according to different monomers within the same protein complex, suggesting the flexibility of their relative orientation. Also, the interaction between T1295-D1 and T1295-D2 (red circle in **Fig. 8A** and **Fig. 8C**) was known from an experimental structure (PDB id: 6E14) showing 100% sequence identity to the target, which was solved before CASP16. Finally, two “interacting” EUs, T1269-D2 and T1269-D3 (magenta circle in **Fig. 8A** and **Fig. 8D**), are connected by a small domain. The boundary between T1269-D2 and T1269-D3 was incorrectly placed in the middle of the small domain due to errors in domain parsing. The observed “interaction” between them only happens within this small domain, and outside this small domain, the two EUs do not interact.

**Figure 8.**
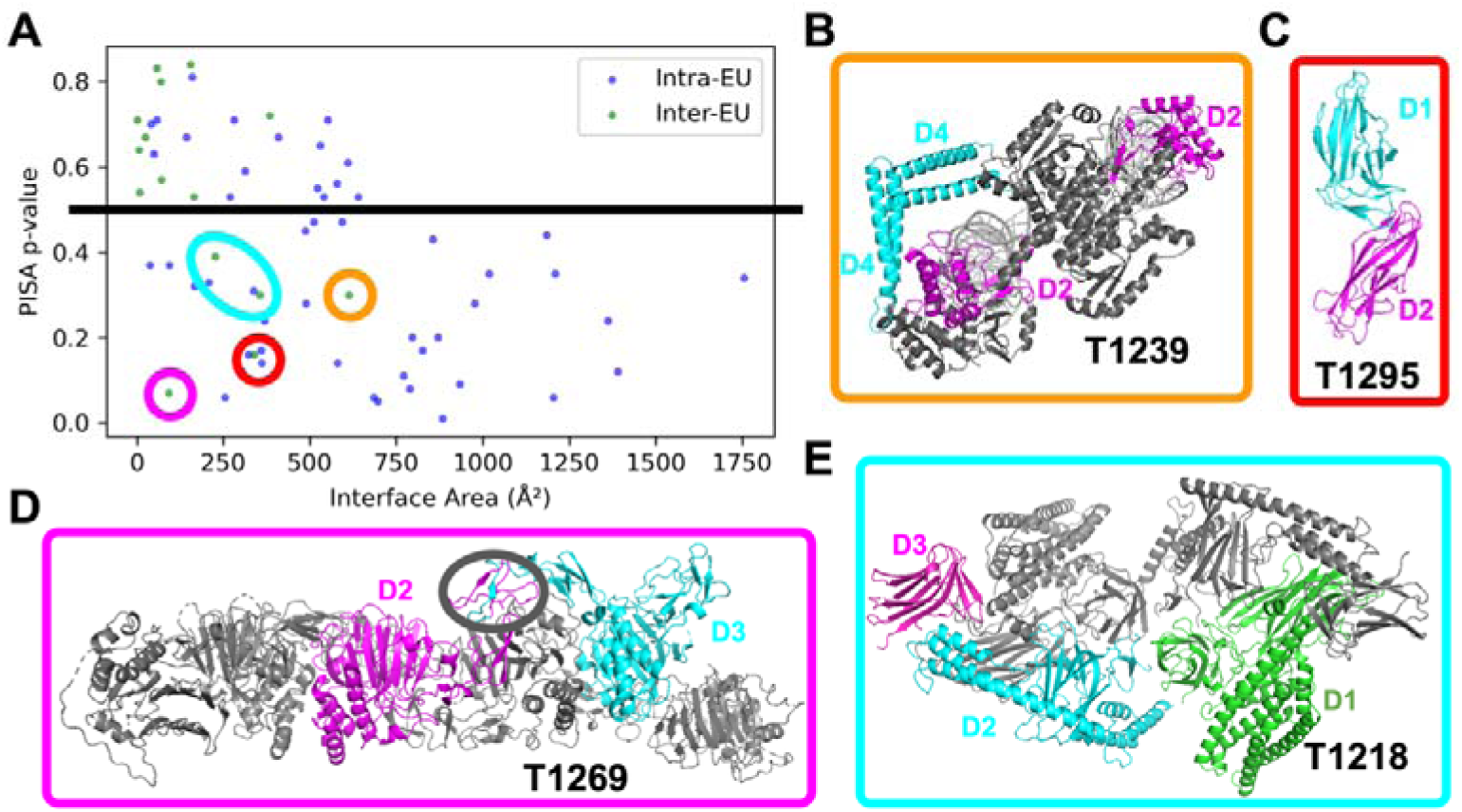
**A)** PISA interface size versus p-value for biological relevance for intra- and inter-EU DDIs. Each dot represents a DDI and inter-EU DDIs showing PISA p-value below 0.5 are marked by circles of different colors, and these colors link these dots to the corresponding structures shown in **B-E. B)** A target contains multiple copies of the same chain adopting different conformations. In one possible conformation, T1239_D2 (magenta on the right) and T1239_D4 (cyan on the top) do not interact; in another conformation, T1239_D2 and T1239_D4 interact. **C)** Two EUs from this target, T1295_D1 and T1295_D2, are interacting, but the 3D structures of these two EUs are available in PDB prior to CASP16. **D)** Two EUs appear to “interact” with each other due to a domain boundary issue. T1269_D2 and T1269_D3 are connected to each other in sequence, and the boundary between them is within a smaller domain (in the black circle). The two EUs do not interact with each other except in this small domain. **E)** Three EUs from T1218 with two biologically meaningful interfaces: between T1218_D1 and T1218_D2, and between T1218_D2 and T1218_D3.

Only the two interfaces formed by T1218-D1 and T1218-D2, and T1218-D2 and T1218-D3 (cyan circle in **Fig. 8A** and **Fig. 8E**) are likely to be biologically meaningful and suitable for DDI prediction and evaluation. Therefore, we reason that the evaluation of inter-EU interaction became unnecessary in this CASP, given that the significant interactions are already within each EU, and they are all correctly predicted by the top predictions (supplemental **Fig. S1**)

## Conclusions and Future Perspectives

Single-domain protein structure prediction is now largely a solved problem, as evidenced by CASP16, where not a single fold was incorrectly predicted. However, specific challenges persist that continue to test the limits of current methods. Larger proteins, as well as viral and eukaryotic proteins, and those with shallow multiple sequence alignments (MSAs), remain more difficult to model accurately. Structural features such as truncated regions, irregular secondary structures, and interchain regions are also prone to prediction errors. These cases highlight important directions for further development within the prediction community.

As in CASP15, the top-performing strategies in CASP16 relied heavily on AlphaFold2 (AF2) and AlphaFold3 (AF3). Our analysis of Phase 2 predictions and the MassiveFold results confirms that sampling more structures has the potential to improve outcomes. Additionally, constructing better MSAs is likely to enhance prediction accuracy. However, due to the limited submissions for the “Model 6” trial, the benefit of improved MSA quality has not yet been rigorously benchmarked. We encourage broader participation in Model 6 in future CASP experiments, as it provides a valuable opportunity to assess MSA quality independently of network architecture.

Overall, performance on monomer targets in CASP16 showed minimal improvement compared to CASP15. This is consistent with the observation that simply increasing AF2 sampling often yields models comparable to or better than those produced by AF3. That said, AF3 demonstrated superior accuracy in confidence estimation, and further optimization following its full release could lead to new breakthroughs in the field.

Community engagement in the additional CASP16 challenges—Phase 0, Phase 2, and Model 6—was notably lower than in the traditional Phase 1 challenge. Nevertheless, we believe these newer challenges are critical for driving future progress. Predicting complex structures without prior stoichiometry information, as in Phase 0, represents an important frontier and should be considered as a core component of future CASP evaluations. Phase 2 aimed to stimulate development in model ranking, but its current design conflates model generation and ranking. A more meaningful approach would be to ask predictors to select their top five models from a fixed set provided by organizers. To encourage participation in Model 6, organizers and future assessors could consider including it in the main performance evaluation, using each group’s best model as the basis for comparison. This approach would incentivize submissions of model 6 by giving participating groups additional opportunities to achieve higher rankings.

The field of protein structure prediction has witnessed remarkable progress in recent years, largely driven by the introduction of deep learning methods. Looking ahead, we hope this manuscript can provide a comprehensive evaluation of the current status of protein structure prediction while also highlighting the remaining challenges that could be focused on to continue advancing the field.

## Methods

### Definition of evaluation units (EUs)

The original EUs were defined according to the CASP15 principles^39^ and further manually examined. We made some adjustments based on the following criteria: 1. Merging EUs. We studied whether any pairs of EUs from a target should be merged based on the Grishin plot—initially proposed by Grishin and colleagues in CASP9^82^. If merging two EUs does not remarkably affect (supplemental **Fig. S14**) the measured prediction accuracy by GDT_TS, we would merge them. 2. Consolidating targets with different conformations. In this CASP, several EUs (T1214-D1, T1228-Ds, T1239-Ds, T1249-D1, T1294-D1) have multiple versions due to different conformations and ligand-induced fit. We evaluated the structural variations between different versions, and when such differences, as measured by RMSD, were above 1□, we used different versions in our score calculation. Otherwise, we used the version 1 designated by the Prediction Center. Different versions of a target were evaluated collectively: model comparisons were conducted against all available conformations, and the highest score obtained from any version was used for the assessment. However, for simplicity and consistency in structural visualizations, only v1 was used in all figures and presentations. 74 EUs were thus defined for Phase 1, and we propagated these definitions to the corresponding targets in Phase 0 and Phase 2, yielding a total of 145 EUs combining all 3 Phases.

### Heatmap Generation and Clustering

The highest GDT_HA from each group for each model was collected. Groups submitting less than 25% of the targets were removed. Ward clustering algorithm was applied to predictors and targets based on performance similarity. Clustering algorithms typically require a definition of distance, and in this case, missing values were assigned 50 for simplicity. The orders in heatmaps reflect the results of clustering and do not imply ranking. The annotations describing predictor strategies and target characteristics were derived from participant abstracts and official CASP classifications, respectively.

### Performance evaluation and ranking

As mentioned in the main text, we included nine scores, i.e., GDT_HA, AL0_P, CAD_AA, GDC_SC, SphGr, lDDT, reLLG_const, QSE, and MolProbity score. These scores evaluate model accuracy across multiple dimensions: global fold correctness, local geometric precision, and suitability as templates for experimental studies and other applications. Here we briefly describe each score. **GDT_HA**^58^: Measures the percentage of correctly aligned residues after structural alignment, emphasizing high-accuracy similarity in structures; **QSE**: Measures the difference between pLDDT^6^ and lDDT^83^, serving as a measure of self-assessment; **reLLG_const**^65^: Evaluates the utility of models in molecular replacement (MR) by estimating the log-likelihood gain, a value in MR that measures how much better the experimental data can be predicted with a model than with a random distribution of the same atoms. This score highlights the usefulness of the model for structural biologists; **SphGr**^84^: Evaluates protein structure models by comparing local spherical regions of the model with the target at multiple scales; **CAD_AA**^85^: Evaluates protein model accuracy at an atomic level by comparing the computed residue contact areas between the model and target; **GDC_SC**^58^: Measures the percentage of correctly aligned sidechain atoms; **AL0_P**^58^: Represents the percentage of residues correctly aligned in the model based on the LGA superimposition; **lDDT**^83,86^: Evaluates model-target local distance differences without structural superposition; **MolProbity score**^55^: a comprehensive measure that evaluates how well the model follows physicochemical rules.

Following the tradition of CASPs, we used Z-scores for performance evaluation and ranking. Prior to Z-score calculations, we identified cases where certain models could not be scored on specific metrics due to intrinsic model limitations. In these instances, missing values were imputed using the lowest observed value across all models for the target in that metric. Z-score calculations were carried out in two rounds. In the first round, models with an initial Z-score below −2 were considered outliers and excluded. Z-scores were then recalculated, and for ranking purposes, any Z-scores below −2 were imputed as −2. This approach ensures that groups employing more experimental or speculative methods, which may occasionally result in poor models, are not disproportionately penalized and can still be recognized for consistently strong performance on other targets.

As in previous CASP assessments, final rankings were based on the sum of Z-scores across all targets, emphasizing overall performance. To address concerns that some models may capture the global fold well while failing in other aspects, the composite scoring scheme was used. We made rankings on both first and best models (the model with the highest Z scores of each group). If the first model was not designated by the group, the model with the lowest model ID was used as the first model.

To assess the robustness of the rankings and evaluate whether performance differences between groups are statistically significant, we performed head-to-head comparisons for the top 25% groups based on the cumulative Z-score-based ranking. In such head-to-head comparisons, we compare the performance of each two groups using only the targets for which both groups submitted predictions. We generated 1,000 bootstrap replicates of these common targets to estimate statistical significance. For each replicate, we compared the groups’ performances based on the summed Z-score over the nine component scores and counted how many times each group outperformed the other across the replicates.

### Sidechain accuracy evaluation

We evaluated sidechain accuracy by two methods. First, we used GDC_sc provided by the organizers. However, GDC_sc is a superimposition-based score and will be largely correlated with the quality of predicted backbones. Therefore, following CASP15 evalutaion’s procedure^30^, we also implemented SCWRL4’s AAA score^87^ to evaluate sidechain quality based on χ-angles, a measure that is independent of the backbone dihedral angles.

### Correlation analysis and Exploratory Factor Analysis (EFA)

To explore underlying relationships between these metrics, we conducted correlation analysis and EFA. The correlation matrix is computed with raw scores across evaluation measures. For EFA, we used the factor_analyzer package in Python^88^. The rotation parameter is ‘promax’, and the method parameter is ‘minres’. Note that EFA does not guarantee a unique solution, so the results for EFA are for exploratory purposes rather than a rigorous derivation of formulas.

### Model 6 evaluation and cross-phase comparison

Model 6 against model 1 evaluation: For each group, in every target where both model 1 and model 6 were available, we computed the ratio of their GDT_HA scores. Each group’s score was calculated as the average ratio across all such targets.

Model 6 against colabfold_baseline model 1 evaluation: For each group, in every target where both this group’s model 6 and colabfold_baseline model 1 were available, we computed the ratio of their GDT_HA scores. Each group’s score was calculated as the average ratio across all such targets.

Cross-phase comparison: To compare performance across phases, we focused on targets common to both phases under comparison: Phase 0 vs. Phase 1 and Phase 1 vs. Phase 2. For each group, we calculated the gain or loss in GDT_HA by computing the difference between the next phase and a previous one on a per-target basis and then averaging the values across all shared targets.

### Quality evaluation

Using the above Performance evaluation and ranking pipeline, QSE ranking was conducted using only the QSE score, and only the first model. To evaluate the potential improvement we can achieve by selecting the best model among the large amount of MassiveFold models, we computed GDT_HA for MassiveFold models using LGA with the following parameters: lga -3 -ie -o1 -sda -d:4 -dgc_sc. The highest score among all MassiveFold GDT_HA was taken. For each target, we compared the best MassiveFold GDT_HA against the scores for the best model, the first model, the first model of the best group, AF3 server model, and colabfold_baseline model. We also recomputed the GDT_HA scores for these CASP models using our version of LGA, and the results were identical to the values provided by the Prediction Center.

### Progress evaluation

The same scoring procedure was applied for CASP15 and CASP16 results. The highest GDT_HA from each group for each target was collected. Groups submitting less than 25% of the targets were removed. Targets that are missing from colabfold_baseline were excluded. Missing values were imputed with the corresponding score from colabfold_baseline, given the assumption that ColabFold is the tool that every predictor can at least use. Average scores were computed using the above data.

To evaluate performance near functional and biologically relevant regions, we defined them as the residues that are within 5 Å distance to any metals or biologically relevant molecules. The residues were taken from the PDB files submitted by experimentalists and mapped to the processed target PDB. The scores were taken from per-residue GDC_all scores from the output of LGA software provided by the organizers. An average score was computed for each target. The scores were then processed in the same way that GDT_HA data is processed. colabfold_baseline was used as a benchmark.

To compare performance between CASP15 and CASP16, we used two statistical methods to select a set of CASP15 targets showing a similar difficulty level distribution to CASP16 targets. We estimated the difficulty of a target by the GDT_HA score of colabFold_baseline’s first model, and we used a Hungarian algorithm to generate a one-to-one mapping between CASP15 and CASP16 targets, while minimizing the difficulty difference between members in the two sets. To introduce randomness into this process, we first used Gaussian Kernel Density Estimation to fit a continuous probability distribution for CASP15 and CASP16 target difficulties (by colabfold_baseline’s GDT_HA scores). Then, a weighted resampling based on the ratio between the two distributions is done to ensure the bootstrap sample from CASP15 resembles CASP16 in target difficulties.

### Inter-EU and Intra-EU evaluation

For EUs that come from the same target, the interaction between each pair of EUs was computed using PISA to obtain inter-EU interaction results. Within each EU, Chainsaw^37^ was first applied to parse them into domains. For each EU having more than one domain, the interaction between each pair of domains was computed using PISA to obtain intra-EU interaction results. PISA was used with default parameters. 5% of the residues were truncated on both ends of the EU or domain sequences before running PISA to minimize the influence by the fact that some EU or domain pairs are neighboring in sequence and are thus covalently connected.

## Author Contributions

RY and JZhang conducted the assessment with supervision from QC and NVG, and with instructions from AK. RDS and JZhou participated in data analysis and provided suggestions for analysis methods. RY and QC prepared the figures and drafted the manuscript. RY, JZhang, QC, AK, and NVG edited the manuscript.

## Acknowledgements

QC is supported by the Endowed Scholars Program in UTSW and I-2095-20220331 from Welch Foundation. NVG is supported by I-1505 from Welch Foundation and 2224128 from National Science Foundation Division of Biological Infrastructure. AK is supported by NIGMS GM100482. RDS is supported by NIGMS GM147367, JZhou is supported by RR190071 from the Cancer Prevention and Research Institute of Texas, and JZhang is supported by NIAID K99AI180984-01A1. We also acknowledge computing resource support from NSF ACCESS award MED240004 and TACC award MCB23014. The authors thank: All CASP organizers, in particular John Moult, for giving us valuable guidance during CASP evaluation; Gabriel Studer for fixing technical issues in OST software, which help us to gain more insights in the definition of monomer evaluation units; Guillaume Brysbaert for providing MassiveFold structures; Arne Elofsson for providing feedback and pointing out several missing models during our evaluation process; Jianlin Cheng for discussions on target T1257-D1; Yuki Kagaya, Daisuke Kihara for target T1207-D1; and Toshiyuki Oda for his feedbacks during the revision stage of our manuscript.

## Data availability and code availability

CASP16’s official website is https://www.predictioncenter.org/casp16/index.cgi. Raw results can be found at https://predictioncenter.org/casp16/results.cgi?tr_type=regular. Processed data, materials, and code are available upon request.

