## supplemental figures for "CASP16 protein monomer structure prediction assessment"

**Figure S1.** Model with highest GDT_HA (green) aligned with target structure (cyan). If there are multiple models with the highest GDT_HA, one model will be randomly selected. Text colored in red indicates it is a challenging target for most of the predictors.

**Figure S1 (continue).** Model with highest GDT_HA (green) aligned with target structure (cyan). If there are multiple models with the highest GDT_HA, one model will be randomly selected. Text colored in red indicates it is a challenging target for most of the predictors.

**Figure S1 (continue).** Model with highest GDT_HA (green) aligned with target structure (cyan). If there are multiple models with the highest GDT_HA, one model will be randomly selected. Text colored in red indicates it is a challenging target for most of the predictors.

**Figure S1 (continue).** Model with highest GDT_HA (green) aligned with target structure (cyan). If there are multiple models with the highest GDT_HA, one model will be randomly selected. Text colored in red indicates it is a challenging target for most of the predictors.

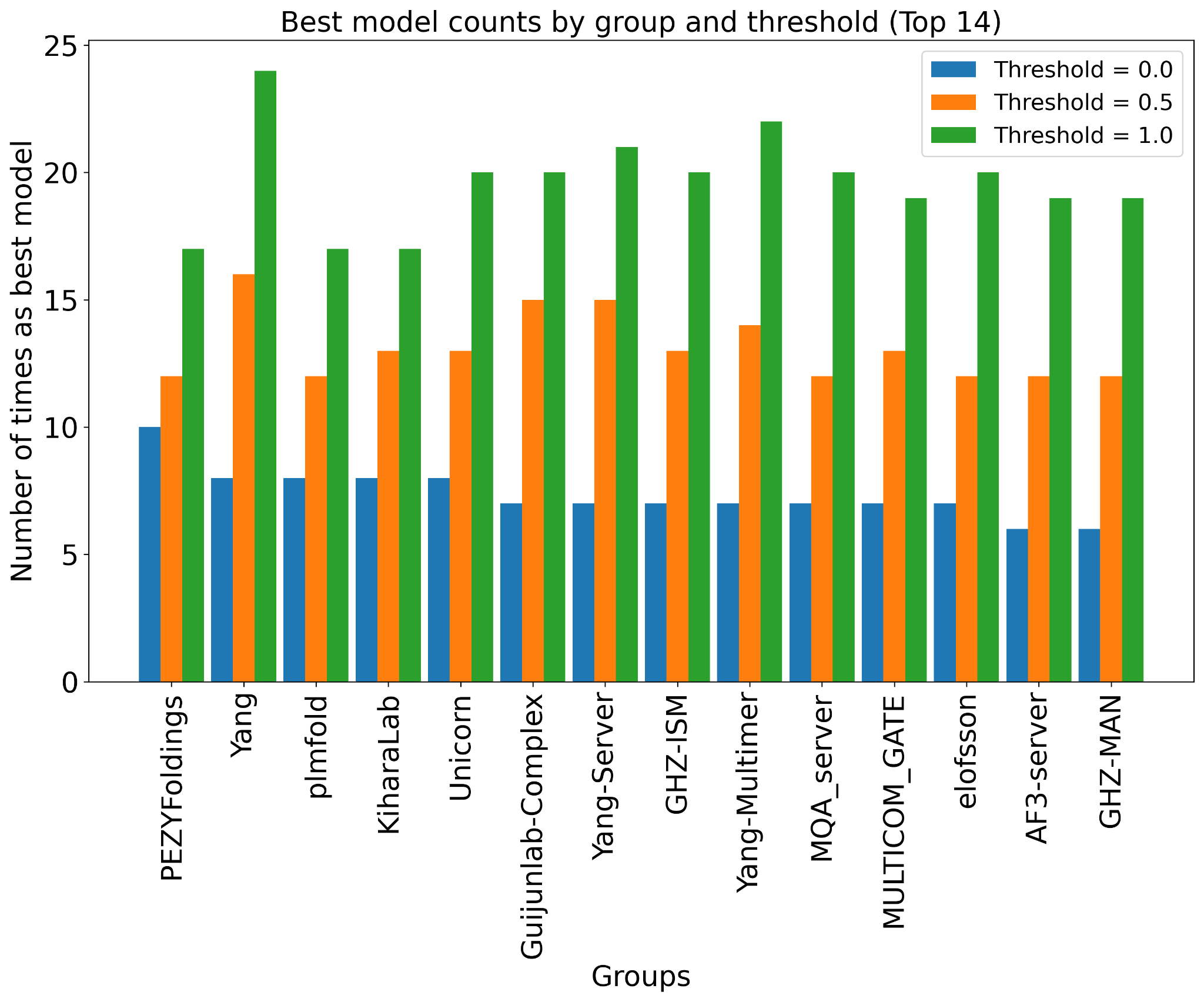

**Figure S2.** The number of best models that one group has. Best models are defined as: GDT_HA >= (highest GDT_HA - threshold).

**Figure S3. (A,B)** Cases where experimental structures might contain poorly resolved regions. **(C)** Predictors’ performance is better on structures with higher experimental resolution.

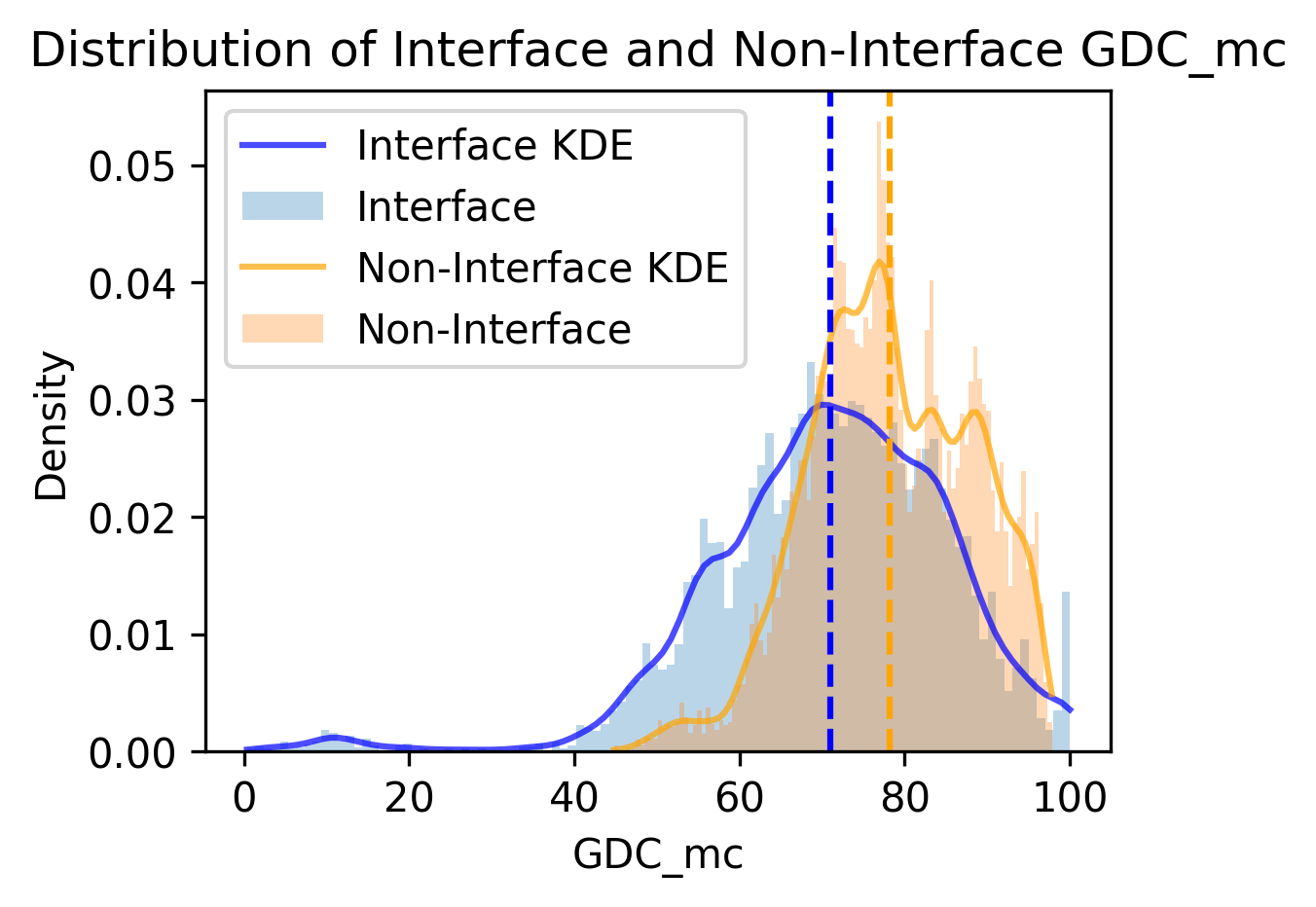

**Figure S4.** Comparison of GDC_mc scores for interface and non-interface residues.

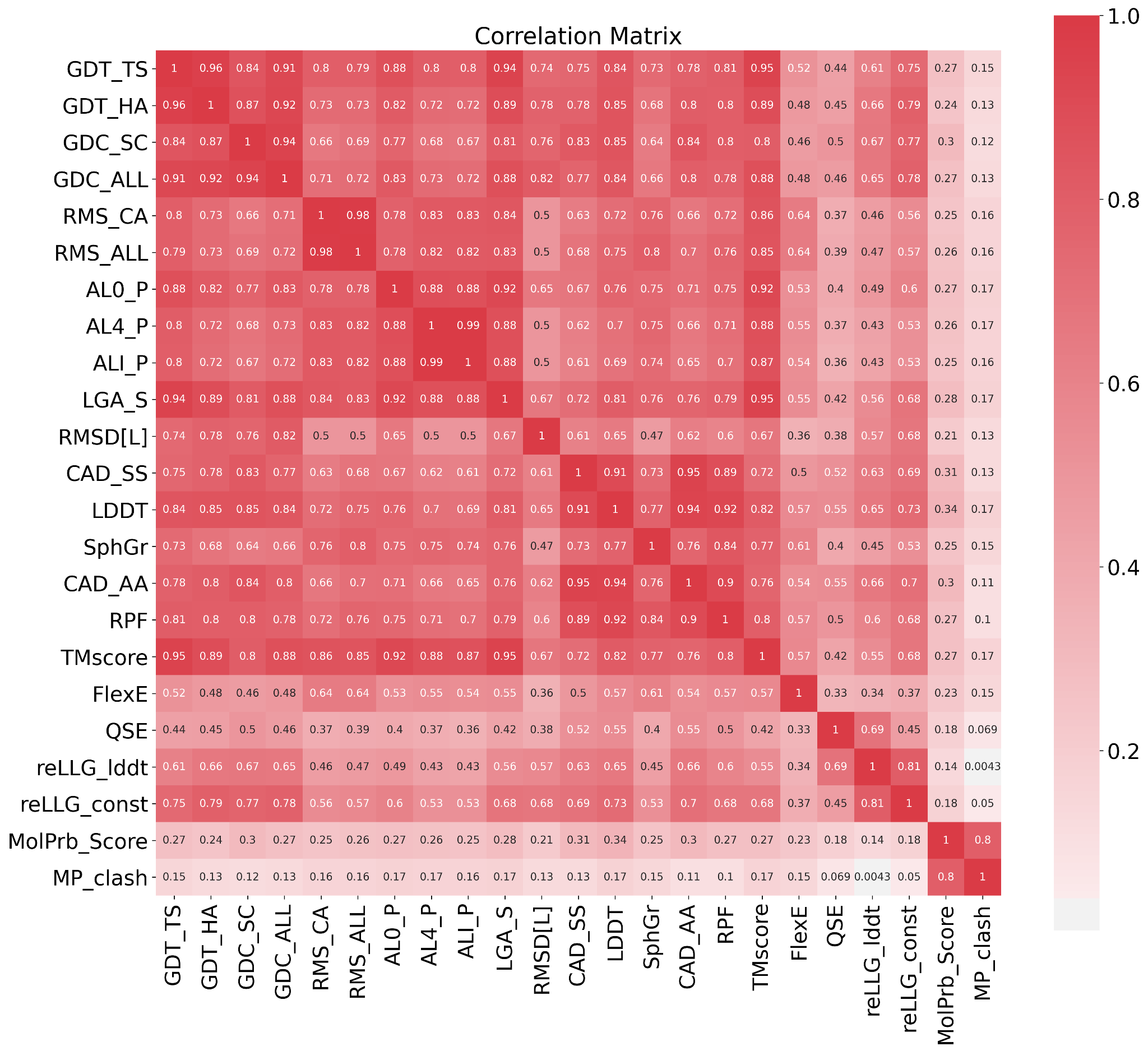

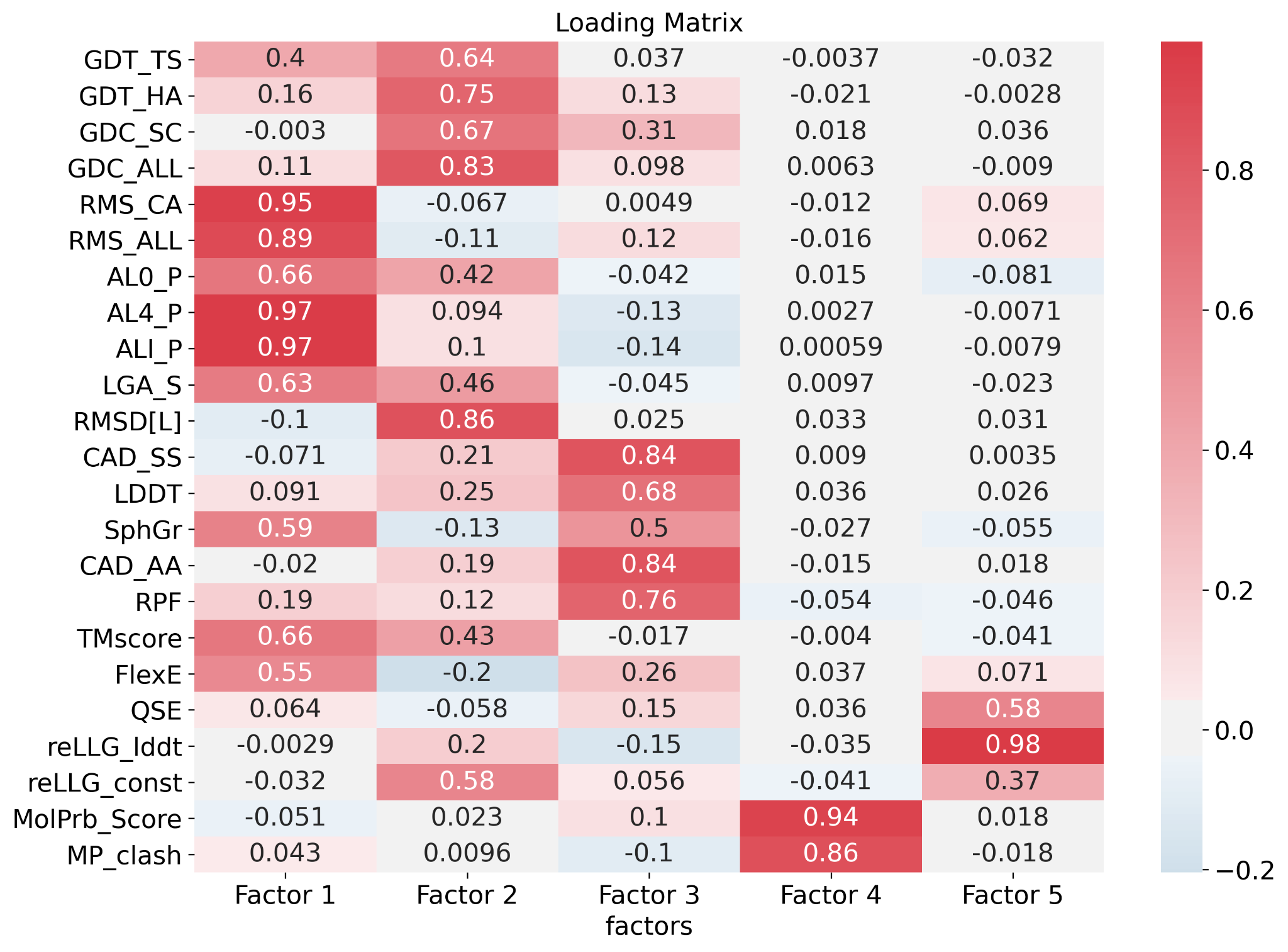

**Figure S5. (A)** Correlation matrix for scores. **(B)** Loading matrix from factor analysis.

**A**
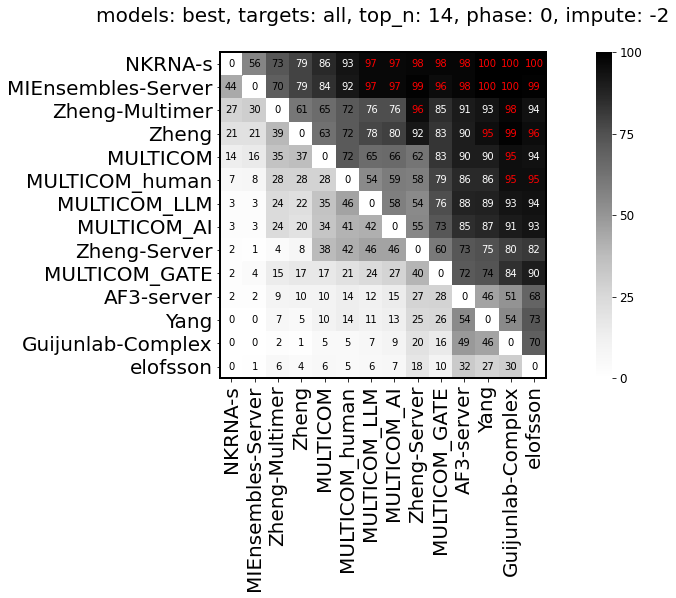

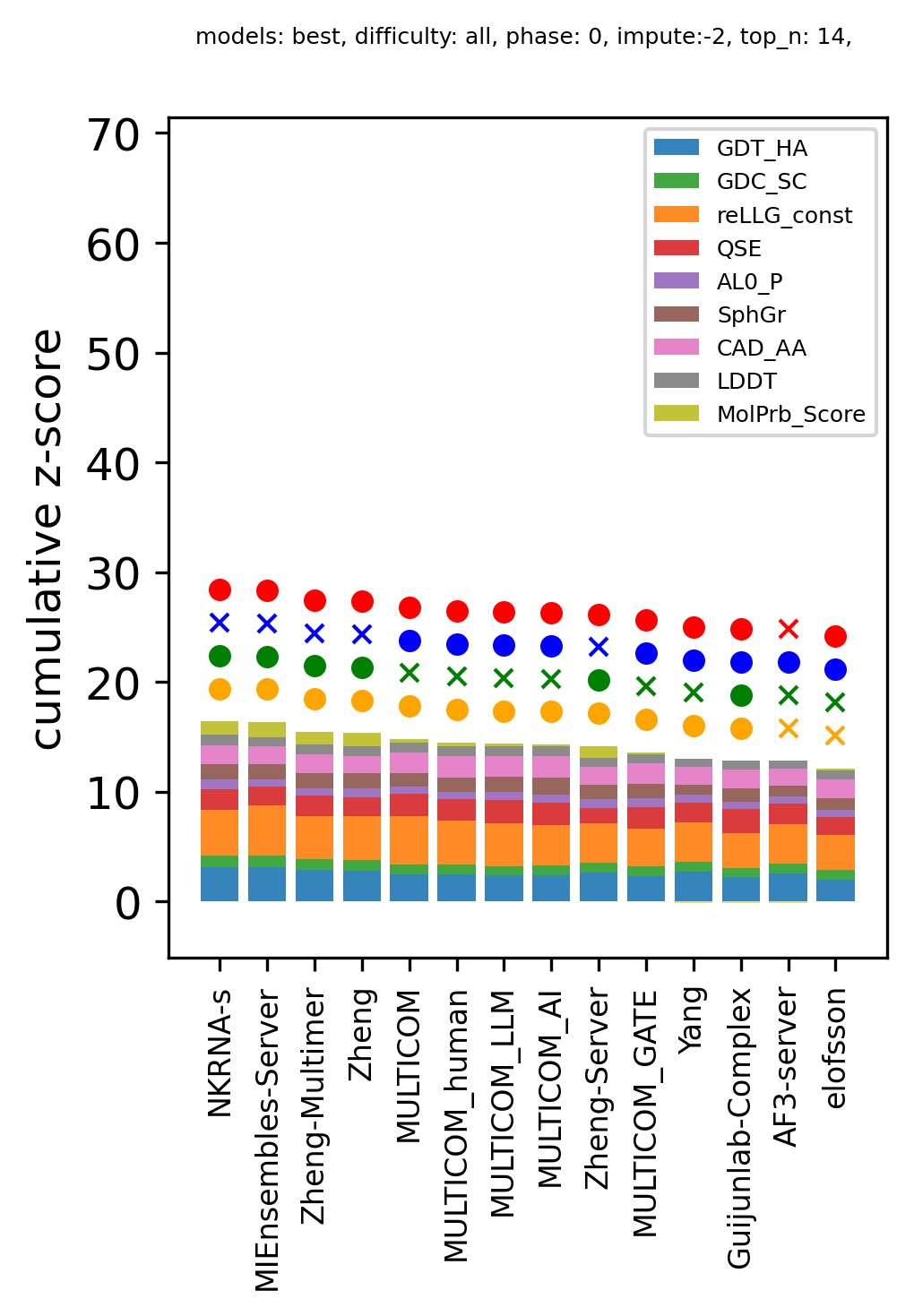

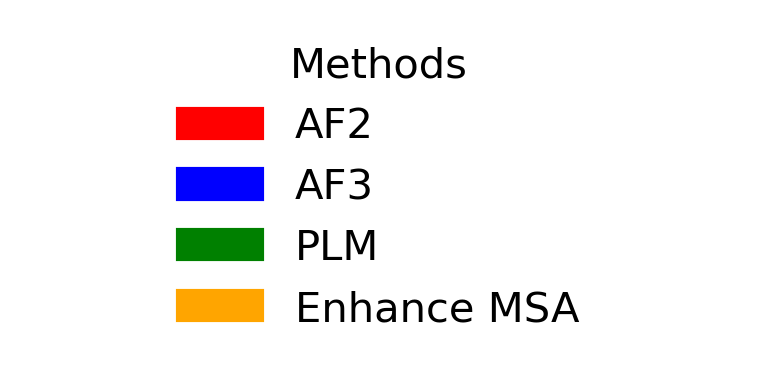

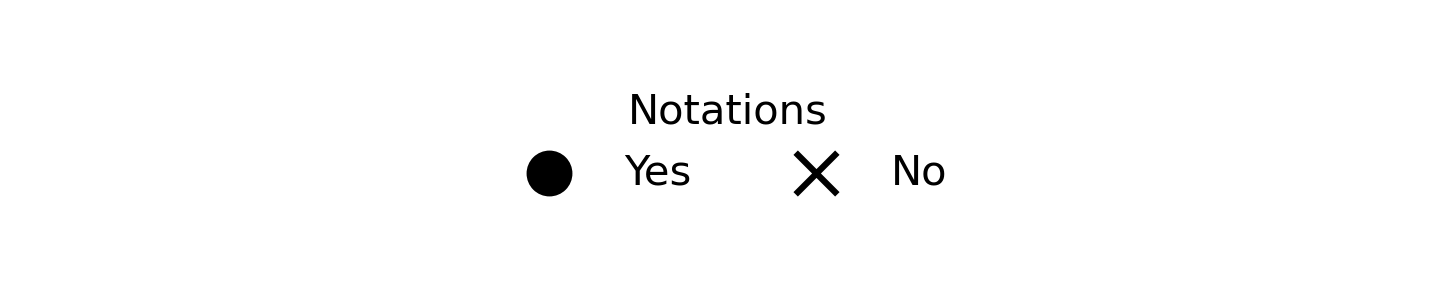

**B**
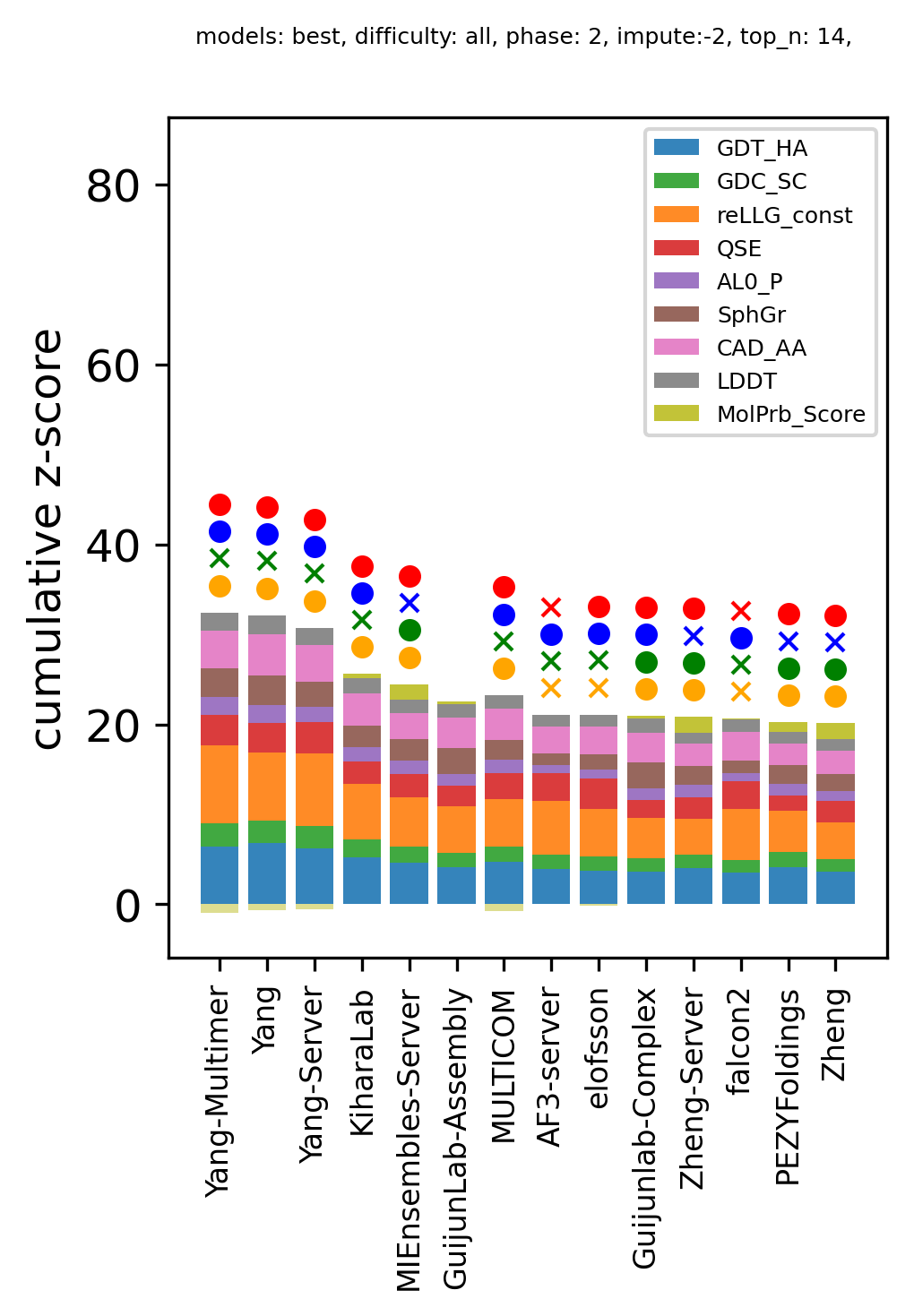

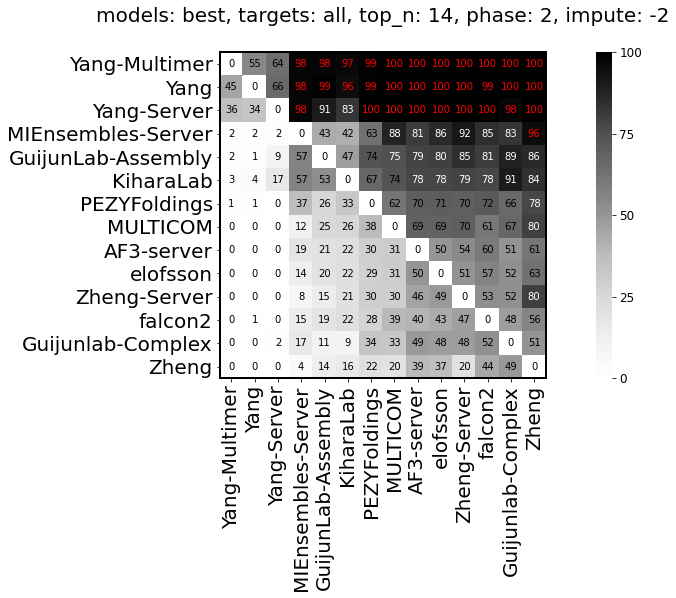

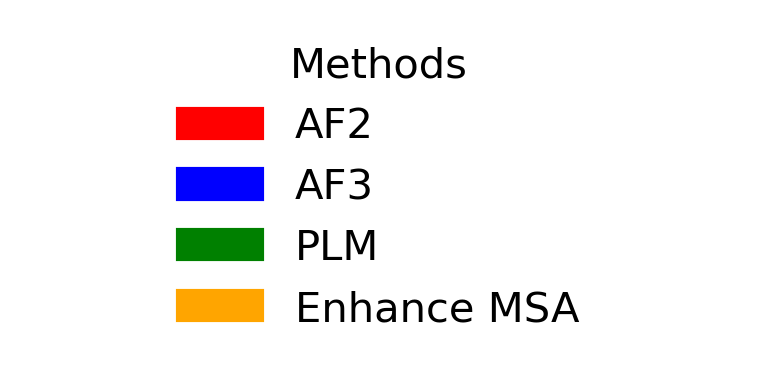

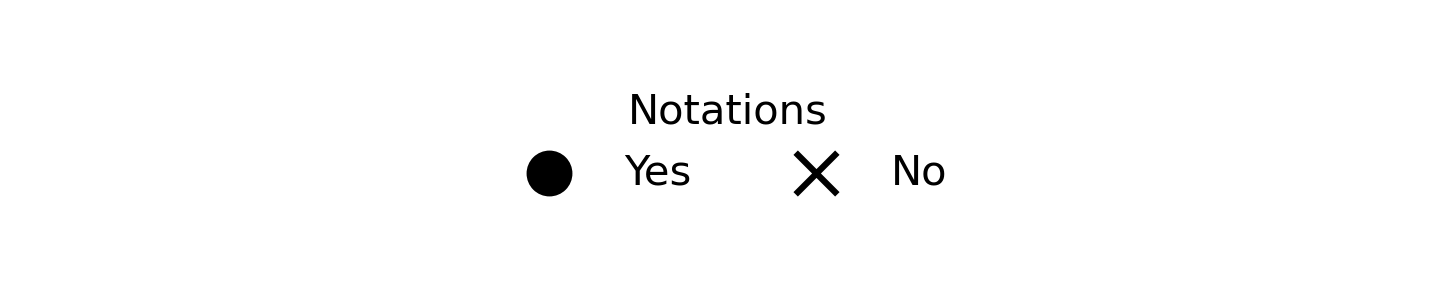

**Figure S6. (A**) ranking for Phase 0 targets (left) and bootstrapping results (right). **(B)** ranking for Phase 2 targets (left) and bootstrapping results (right).

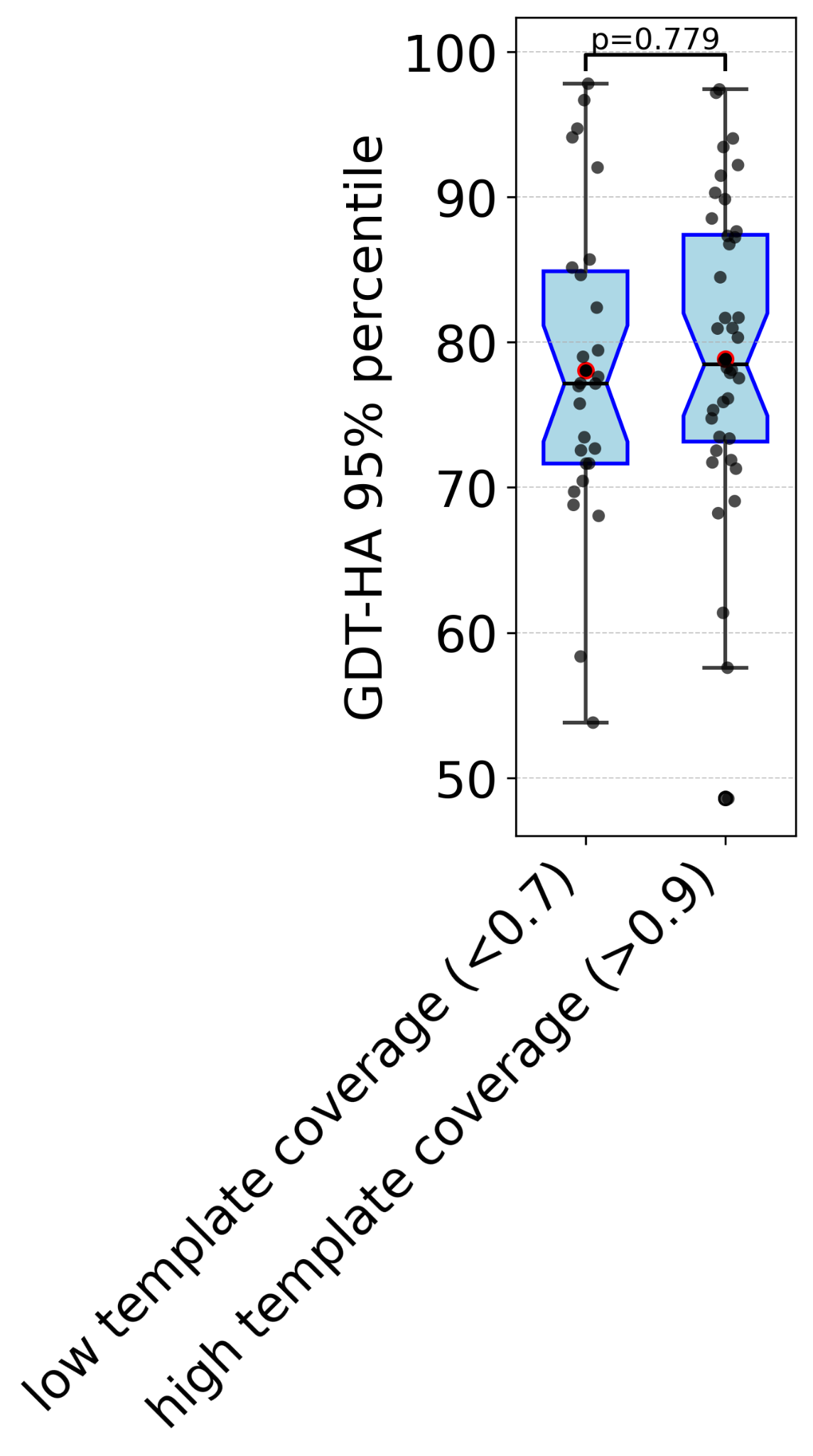

**Figure S7.** Template coverage has no significant effect on prediction quality.

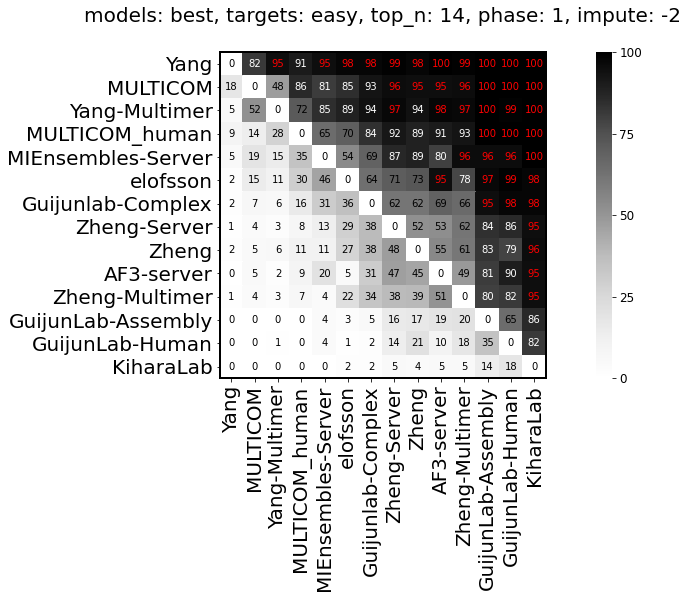

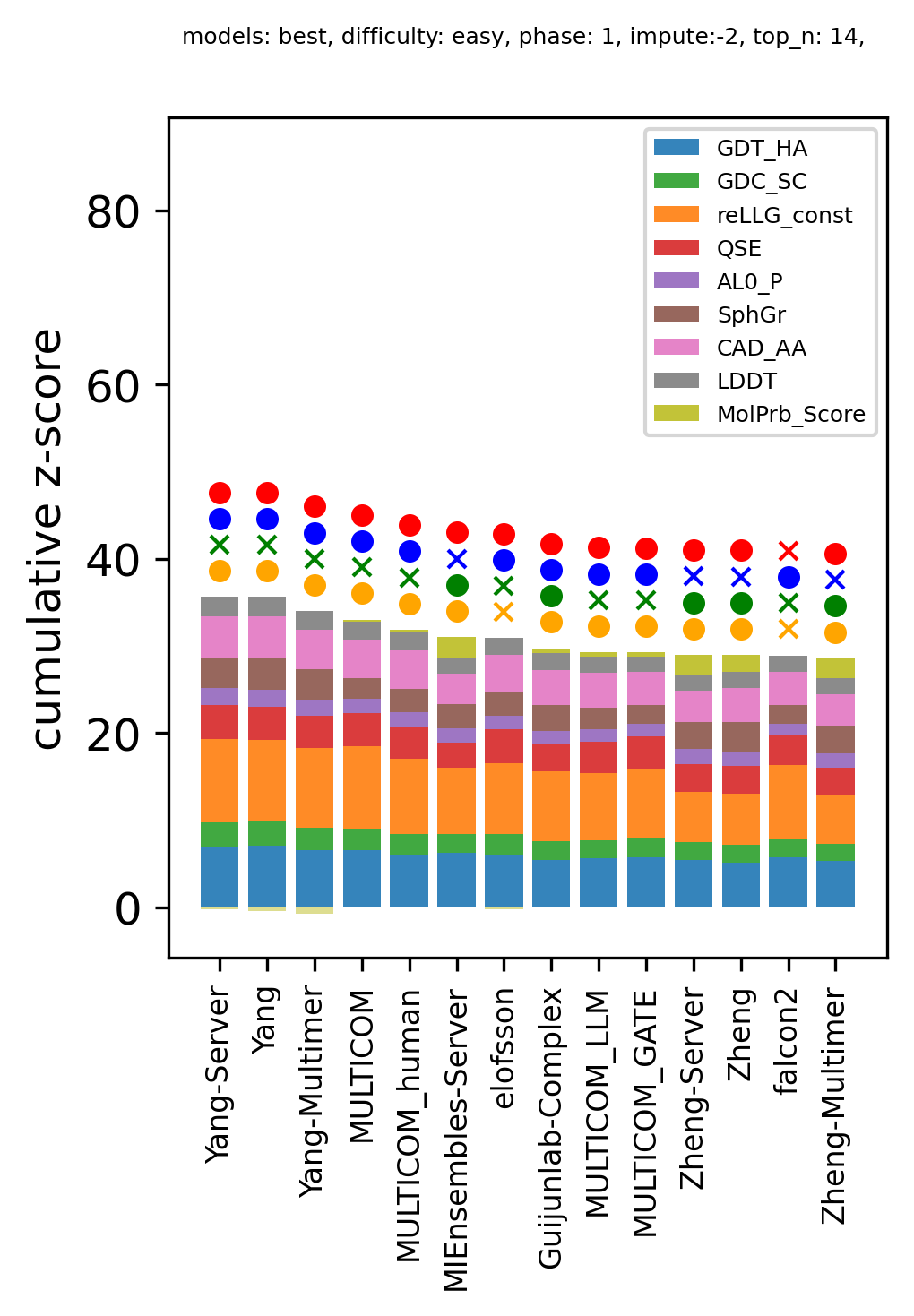

**A**

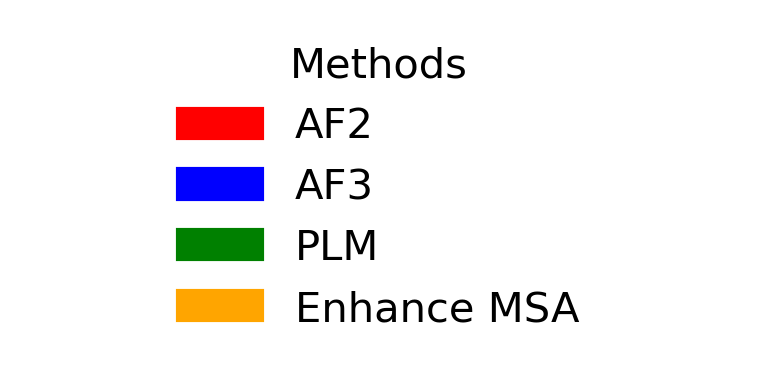

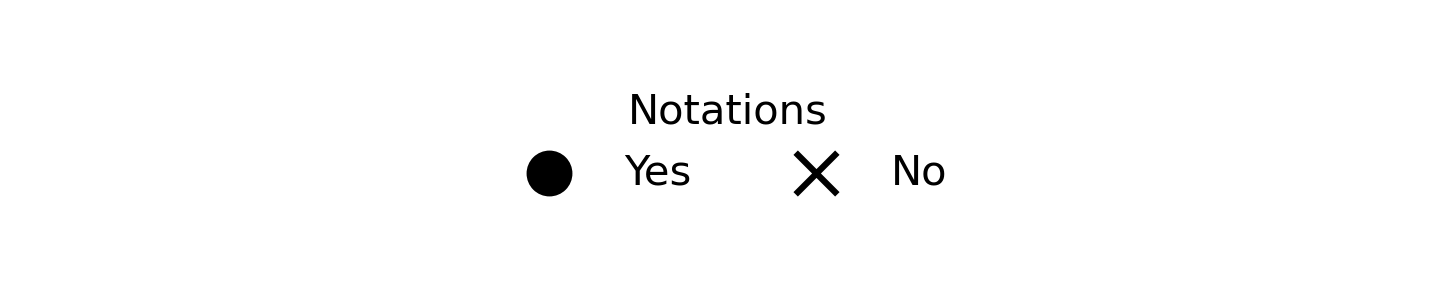

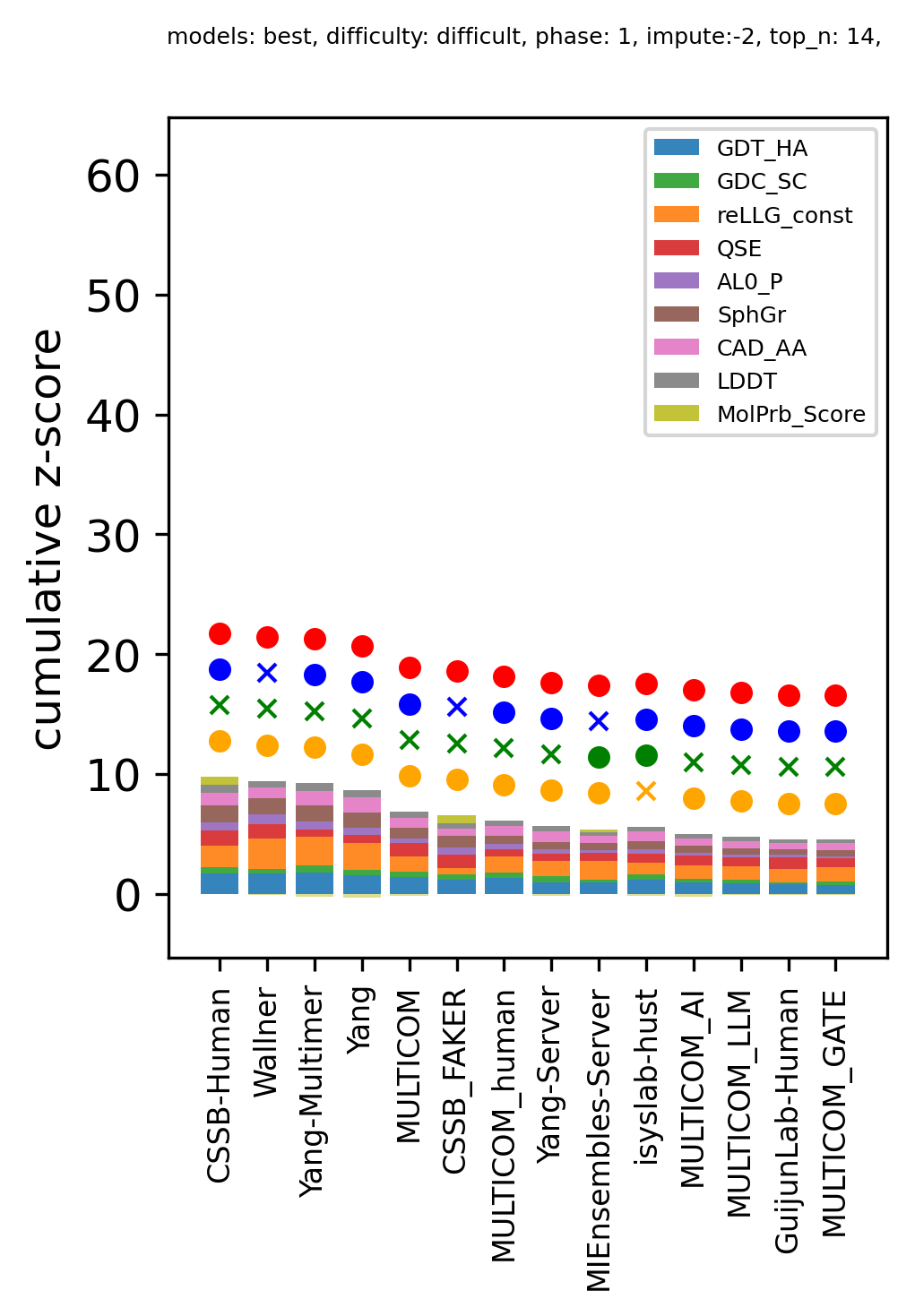

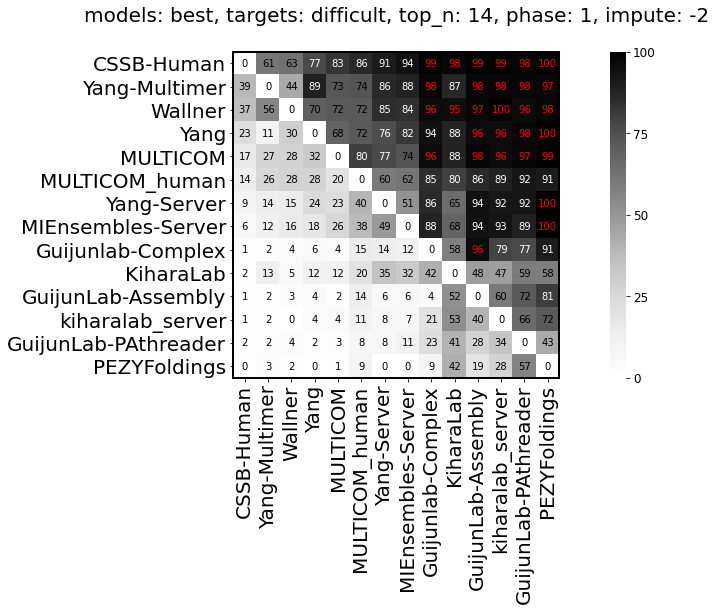

**B**

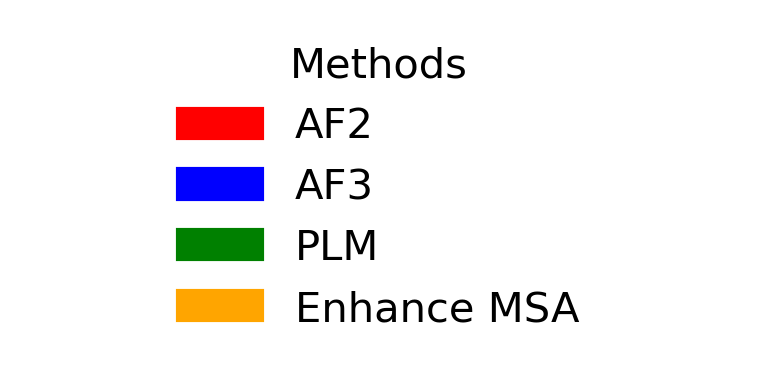

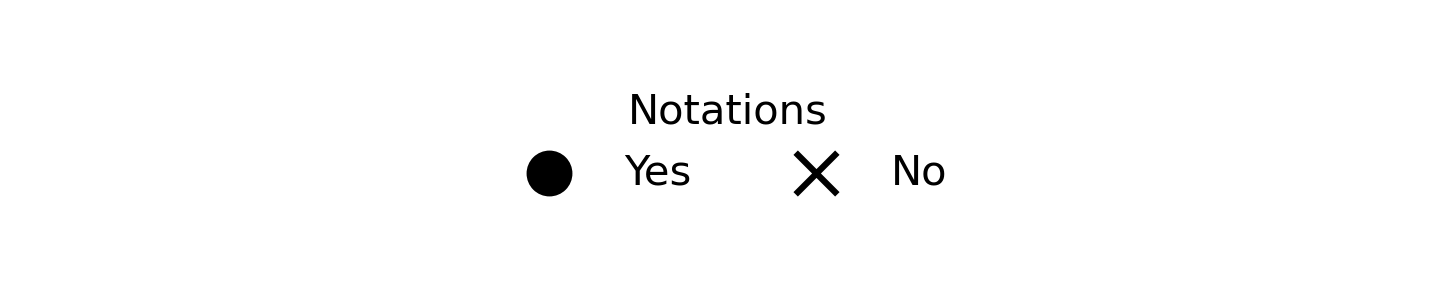

**Figure S8. (A**) ranking for easy targets (left) from Phase 1 and bootstrapping results (right). **(B)** ranking for difficult targets (left) from Phase 1 and bootstrapping results (right).

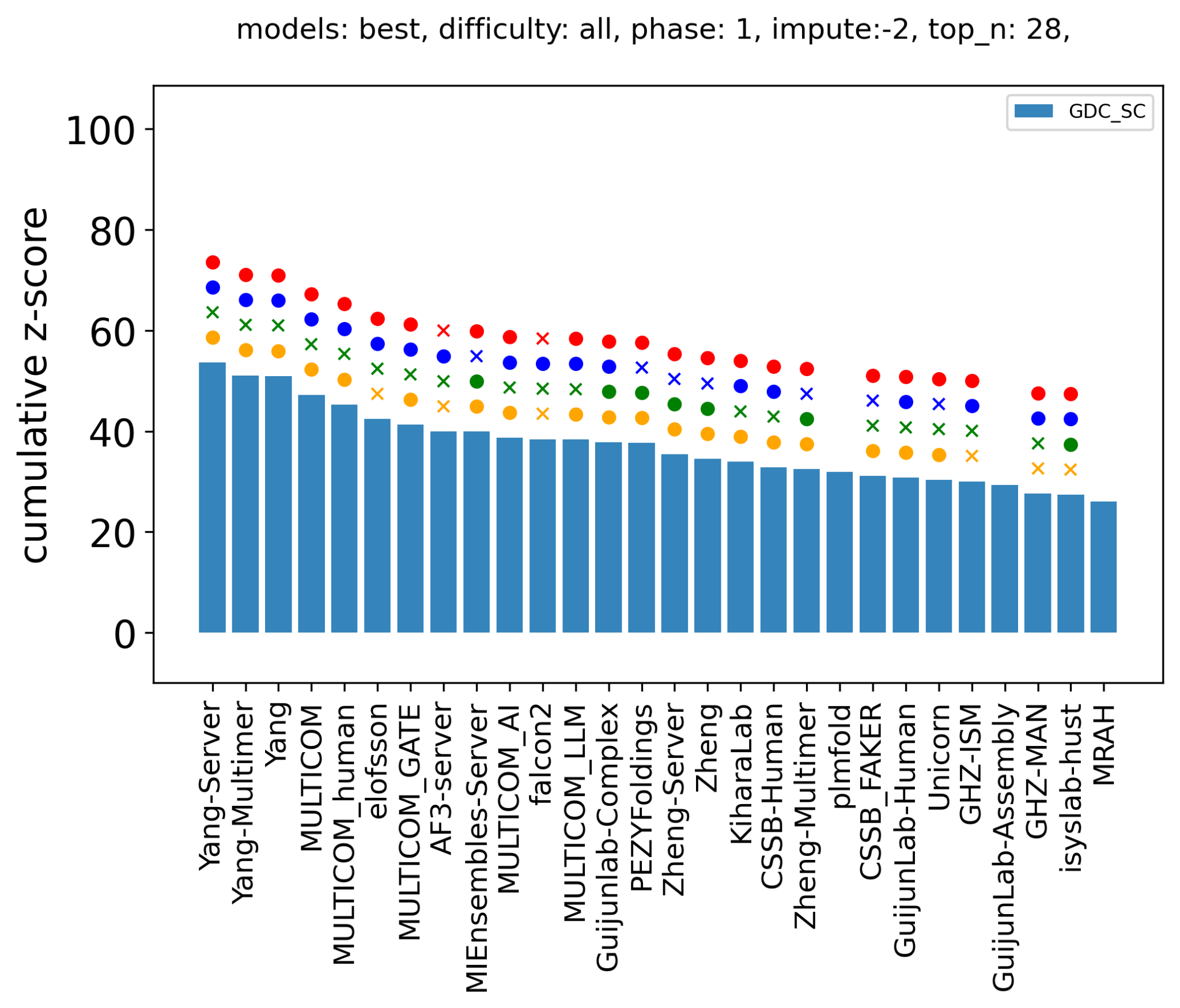

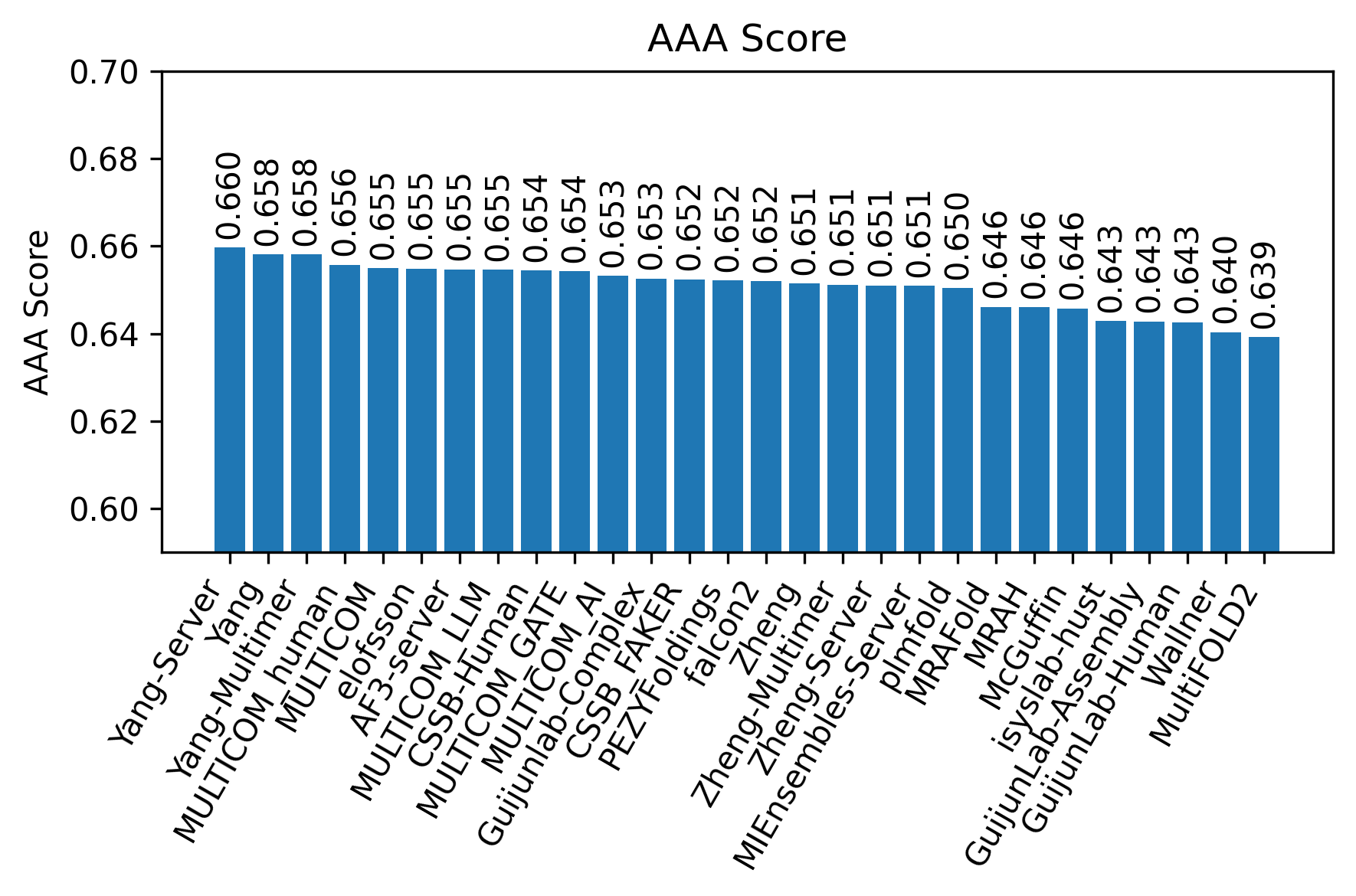

**Figure S9.** Sidechain prediction accuracy of different groups measured by mean GDC_sc scores (top) and mean AAA scores (bottom) for Phase 1 targets.

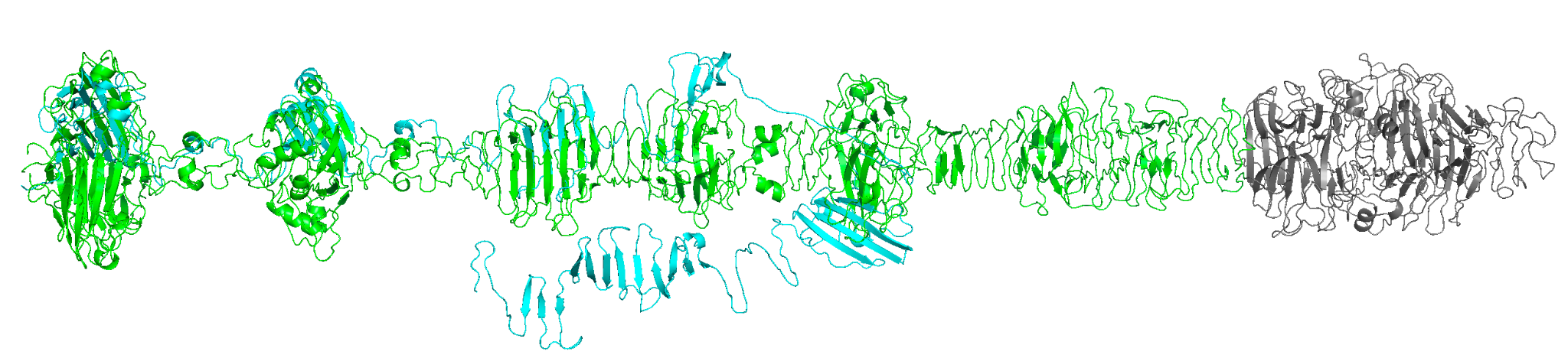

**Figure S10.** A predicted structure by AF3 server for target T1257-D1 (cyan) in its oligomeric target context (green).

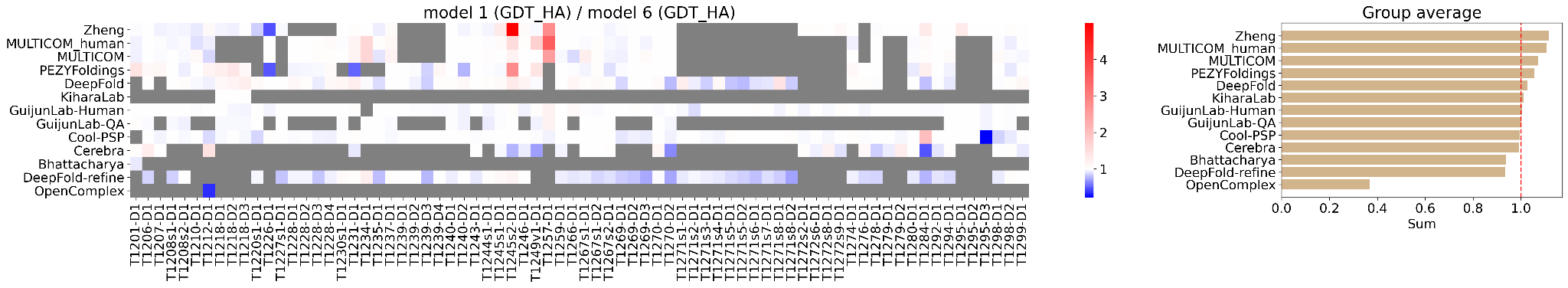

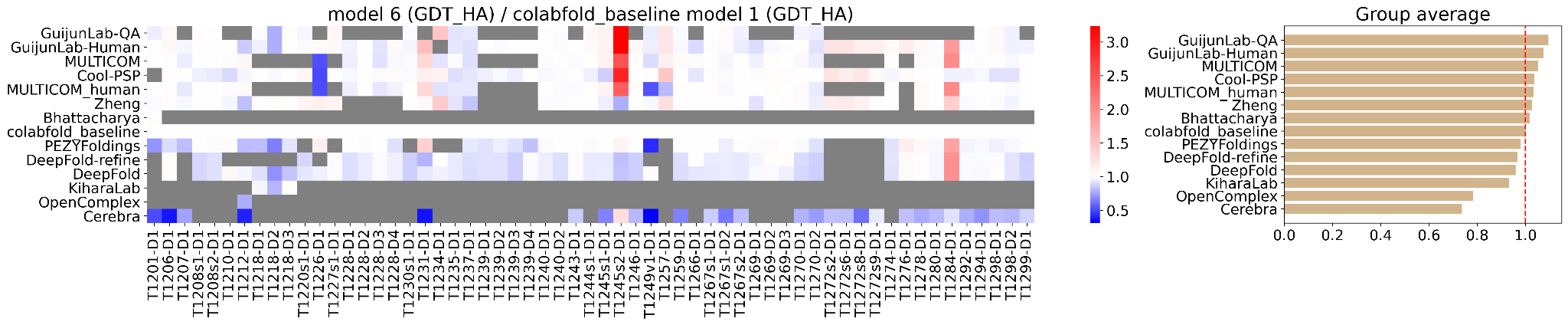

**Figure S11.** Performance evaluation for model 6. Grey indicates N/A. Top: Left: The ratio between model 1 GDT_HA and model 6 GDT_HA. Right: Average of over targets (excluding N/A). Bottom: Left: The ratio between model 6 GDT_HA and colabfold_baseline model 1 GDT_HA. Right: Average over targets (excluding N/A).

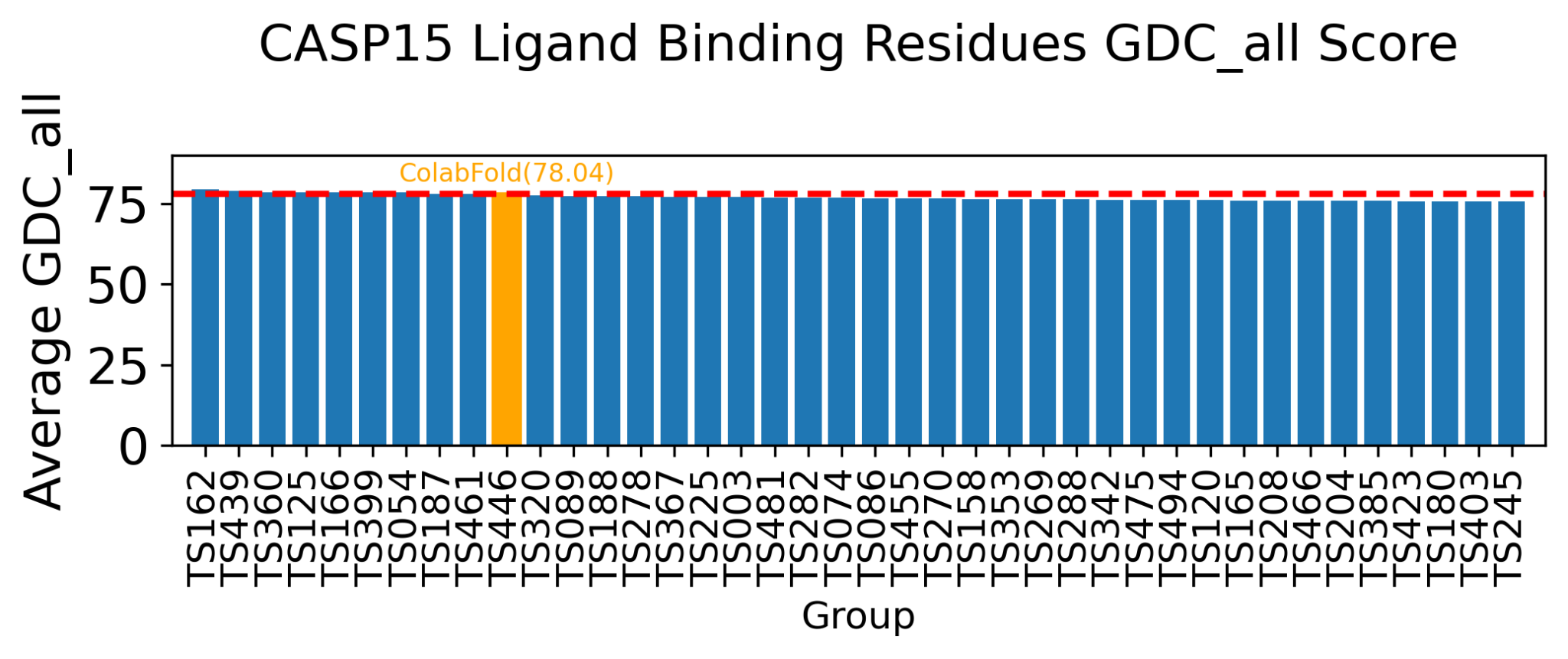

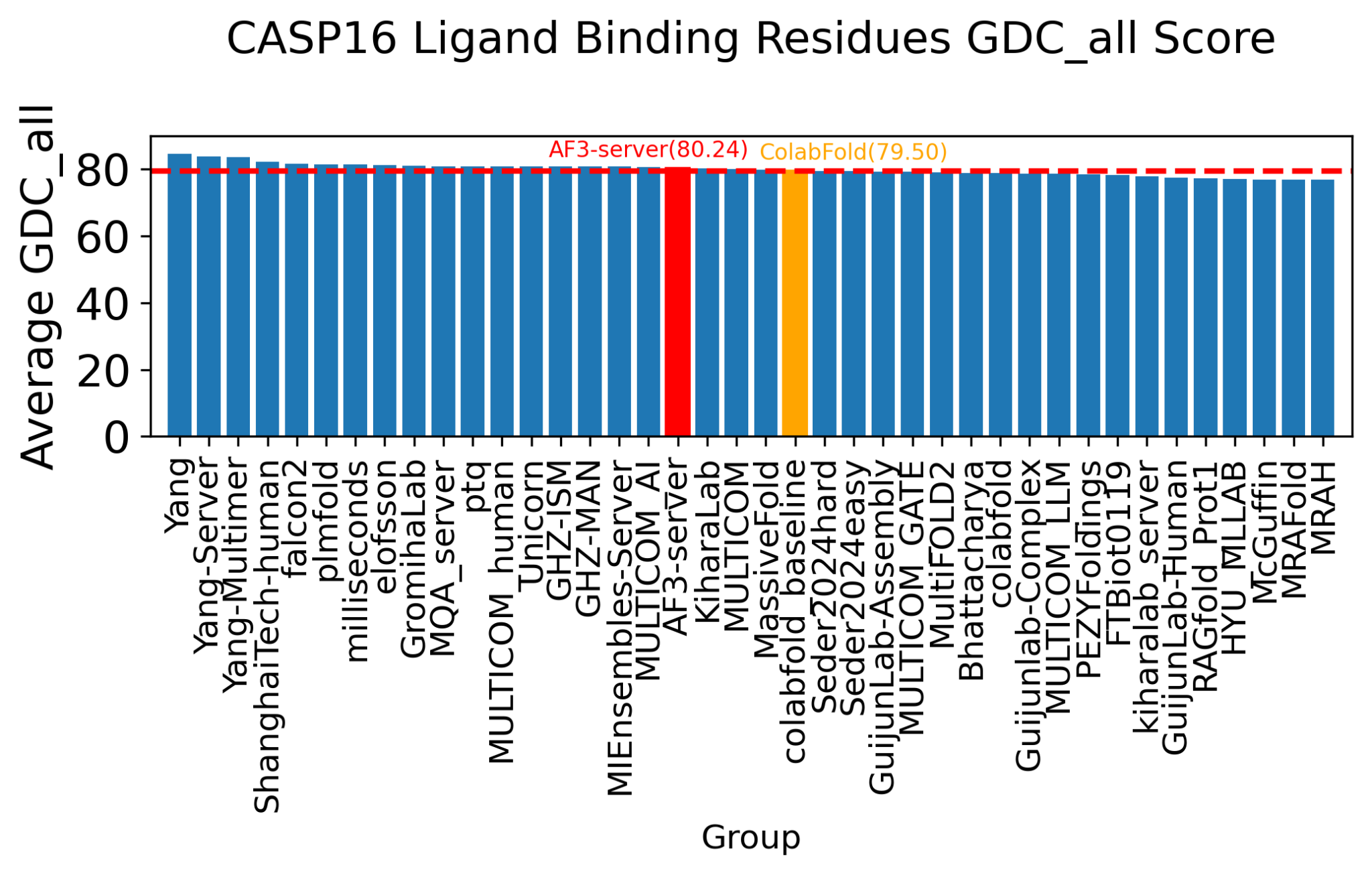

**Figure S12.** Comparison of GDC_all scores extracted from residues that are within 5 Å distance of a biologically meaningful ligand. Top: CASP15; Bottom: CASP16.

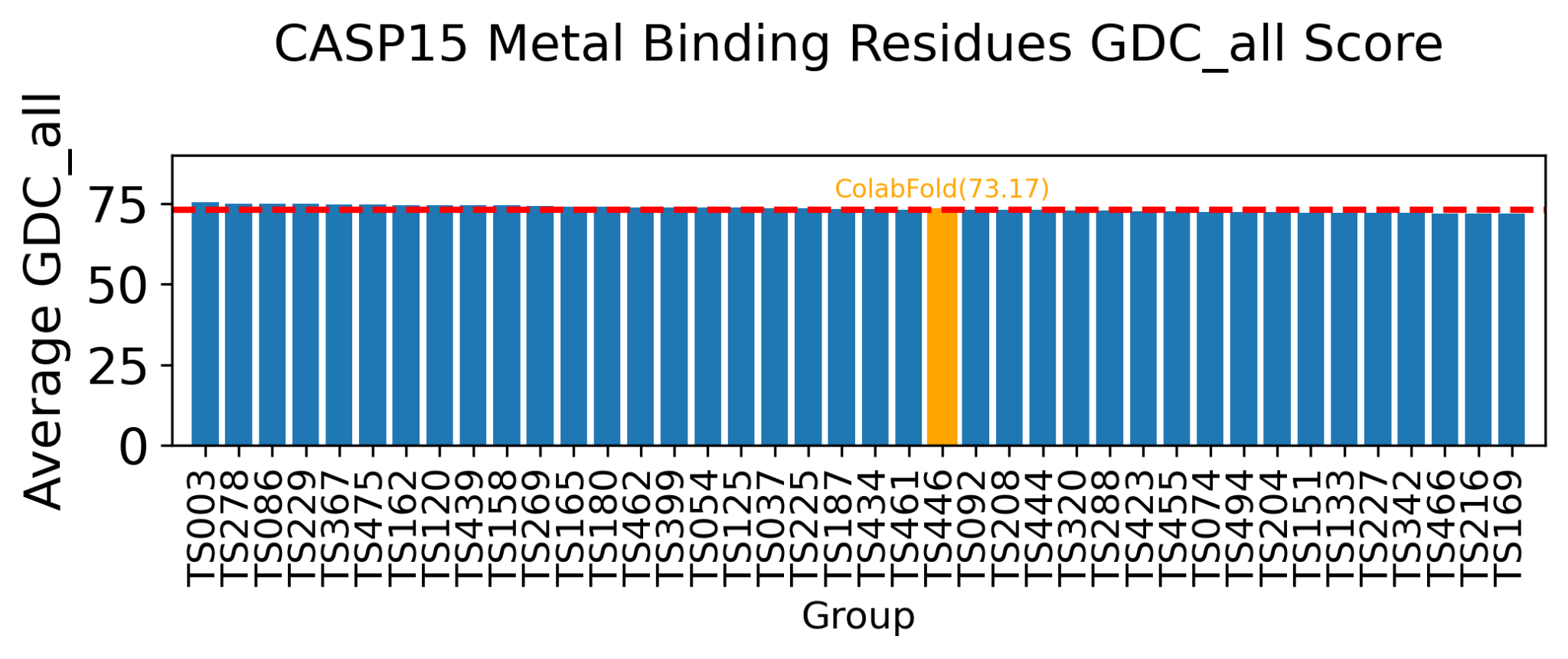

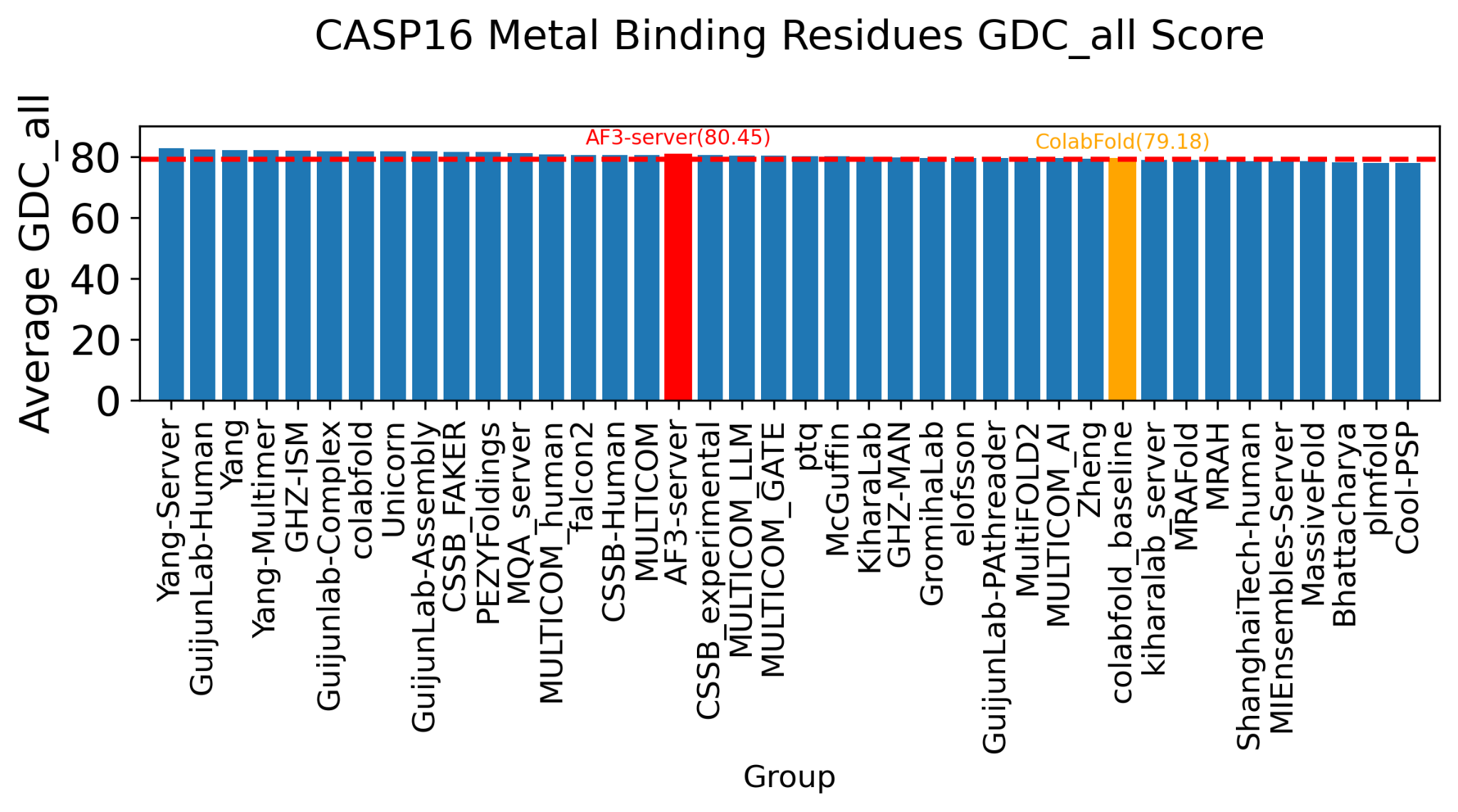

**Figure S13.** Comparison of GDC_all scores extracted from residues that are within 5 Å distance of a metal ion with a charge of +2 or higher. Top: CASP15; Bottom: CASP16.

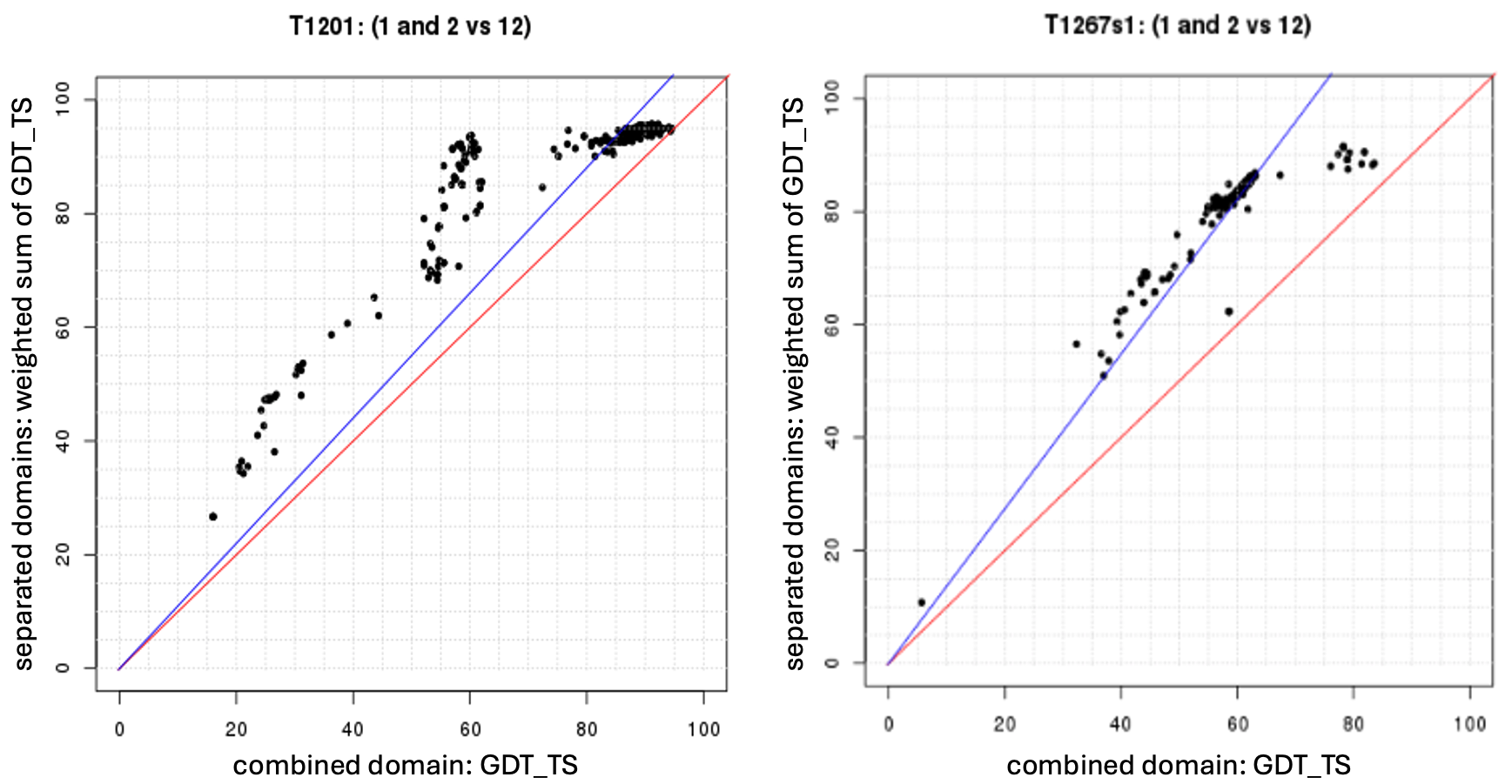

**Figure S14. The Grishin plot is used to determine whether two domains should be merged into a single evaluation unit (EU).** Each plot displays the accuracy of models for a given target. The x-axis represents the GDT_TS when the two domains are evaluated together, while the y-axis shows the weighted (by domain size) sum of GDT_TS scores when the domains are evaluated separately. The red line indicates the identity line (y = x), and the blue line represents a linear regression fit to the observed values (black dots), constrained to pass through the origin. If the angle between the red and blue lines does not exceed 6° (e.g., left panel), the two domains are merged into one EU. Otherwise (e.g., right panel), the domains are treated as separate EUs.
